# Viral Microglia Reprogramming Clears Oligomeric Neurotoxic Debris

**DOI:** 10.64898/2026.04.06.716590

**Authors:** Griffin P. Carter, Zachary P. McKay, Mark A. Katz, Lisbeth Disla, Dasean T. Nardone-White, Derek Southwell, Michael C. Brown, Matthias Gromeier

## Abstract

Owing to pivotal roles in CNS debris clearance and homeostasis, microglia are central targets for the therapy of neurodegenerative diseases. Intricate proximity to neurons, the inherent danger of neuroimmune toxicity, and intrinsically high plasticity and adaptability, impose high hurdles on microglia modulation. Attenuated viruses are being tested extensively against CNS malignancies (i.e., cancer virotherapy); yet, aside from viral vector-mediated payload delivery, virotherapy for non-neoplastic CNS disease remains unexplored. Here we report disseminated targeting of microglia with the highly attenuated polio:rhinovirus chimera, PVSRIPO, that culminated in profound, durable microglia reprogramming. This phenotype, rooted in extended cytoplasmic viral (v)RNA replication, was non-cytopathogenic and did not yield virus progeny or dissemination. vRNA replication in microglia triggered selective interferon (IFN) regulatory factor (IRF) 3/IRF7 transcriptional programs in the relative absence of NFκB-driven proinflammatory cytokine responses and elicited robust phagocytosis of both tumor cells and amyloid-beta. Targeting of microglia with PVSRIPO mediated immunotherapy in a mouse glioma model and the clearance of oligomeric amyloid-beta deposits in an injectable model of neurotoxic amyloid accumulation. This work identifies attenuated virotherapy as an approach to safely and effectively invigorate microglia function in immune surveillance and neurotoxic debris clearance.

## Introduction

Microglia, the tissue-resident phagocytes of CNS parenchyma, stem from erythromyeloid progenitors in the yolk sac that migrate to their destination tissue during embryonic development (Alliot et al., 1999; Ginhoux et al., 2010; Kierdorf et al., 2013). Confined to the CNS, microglia self-sustain via local proliferation, without hematogenous repopulation, throughout life (Ajami et al., 2007). Omnipresence, longevity, and self-renewal align with pivotal roles in CNS homeostasis; development and sculpting of synaptic connections; and defense against trauma and infection. Microglia actively shape malignant, neuro-degenerative, and autoimmune CNS pathologies (Prinz et al., 2019); directing them towards neuro-protective roles is hindered by a delicate anatomical context, phenotypic heterogeneity and plasticity, and intractable functional complexity.

Microglia (and border-associated macrophages, or ‘BaM’) are bulwarks against encephalitogenic RNA (*corona*, *entero*, *flavi*, *paramyxo*, *rhabdo*, *toga*) viruses. Beyond the ubiquitous cytoplasmic vRNA sensors retinoic acid-inducible-I (RIG-I) and MDA5, they are the CNS’ main site for endolysosomal vRNA-sensing toll-like receptor (TLR) 3,7,8 expression (Michaelis et al., 2019). In this study we report striking sustained and disseminated microglia immune reprogramming through non-cytopathogenic infection with polio:rhinovirus recombinant PVSRIPO, the poliovirus type 1 (Sabin) vaccine (PV1S) under control of the human rhinovirus type 2 (HRV2) internal ribosomal entry site (IRES) (Gromeier et al., 1996; Gromeier et al., 1999).

Wild type poliovirus/PV1S naturally target macrophages/dendritic cells (DCs) (Shen et al., 2017): the poliovirus receptor CD155 is constitutively expressed in the mononuclear phagocytic system (MPS). Viral 2A protease-directed cleavage of eukaryotic initiation factor (eIF) 4G (Etchison et al., 1982), the central translation initiation scaffold, leads to lethal host protein synthesis shut-off, blocks host antiviral type-I IFN (IFN-I) defenses, and enables wild type poliovirus/PV1S propagation (Wahid et al., 2005). In contrast, because of encumbered translation of incoming vRNA via the HRV2 IRES (Dobrikov et al., 2022), PVSRIPO cannot direct eIF4G cleavage, block host protein synthesis, intercept IFN-I, proliferate in-, or damage MPS cells (Brown et al., 2017; Brown et al., 2021; Dobrikov et al., 2022; Dobrikov et al., 2023; Ludwig et al., 2025; Mosaheb et al., 2020). Despite these deficits in subduing host cells, PVSRIPO retains the enterovirus strategy of co-opting phosphatidylinosine-4-kinase for forming vRNA replication organelles (Hsu et al., 2010), generating dsRNA intermediates (Brown et al., 2021; Dobrikov et al., 2023). Uniquely amongst RNA viruses, picornavirus RNAs are mainly sensed by MDA5 (Kato et al., 2006; Loo et al., 2008) because of a genome-linked protein that shields vRNAs from RIG-I sensing of 5’-triphosphate (Hornung et al., 2006), and due to an unorthodox host cell entry mode that bypasses the endolysosomal TLRs (Brandenburg et al., 2007). PVSRIPO is under investigation as an intratumor immunotherapy against recurrent glioblastoma in adult patients (Desjardins et al., 2018) and high-grade glioma in pediatric patients (Thompson et al., 2023), demonstrating safety and feasibility of intracranial administration.

Here we report that PVSRIPO infects primary human and murine microglia and directs non-cytopathogenic vRNA replication in the CNS microglial compartment *in vivo*, associated with disseminated changes in morphology. Anti-tumor effects of PVSRIPO in a mouse glioma model relied on microglia/the CNS-resident phagocytic system. Transcriptomic analyses in human *ex vivo* tissue slices showed PVSRIPO vRNA replication in microglia to elicit immune reprogramming phenotypes with select induction of IRF3/IRF7 transcriptional programs. This was associated with robust phagocytic activity against glioma cells and amyloid-beta (1-42) (Aβ), and the induction of markers of immune surveillance functions. Accordingly, we observed clearance and induced meningeal lymphatic uptake of parenchymal oligomeric debris in the Aβ HiLyte intracerebral injection model upon single intracerebral inoculation of PVSRIPO. Our findings demonstrate durable PVSRIPO-directed phenotypic adaptation of microglia towards phagocytic, antigen cross presentation, debris clearance and immune surveillance functions in the brain.

## Results

### Microglia are essential for PVSRIPO-triggered glioma immune surveillance

Clues about microglia’s central role in PVSRIPO immunotherapy stem from mechanistic studies in GBM patient (undissociated) *ex vivo* tissue slices, which revealed microglia/macrophages as the main host targets (Brown et al., 2021). Also, blinded neuropathologist review of glioma-bearing mouse brains identified tissue responses to PVSRIPO dominated by disseminated microglia reactions and proliferation (Yang et al., 2023). To corroborate microglia’s role in PVSRIPO immunotherapy, we used colony stimulating factor 1 receptor (CSF-1R) inhibitor (Plx5622)-mediated phagocyte depletion (**Fig. 1A**). A Plx5622-containing diet reduced microglia in CNS parenchyma (per Tmem119 staining of non-tumor bearing brain) by ∼95% vs. sham-fed animals (72 and 120h on diet), as reported previously (Spangenberg et al., 2019).

**Figure 1.**
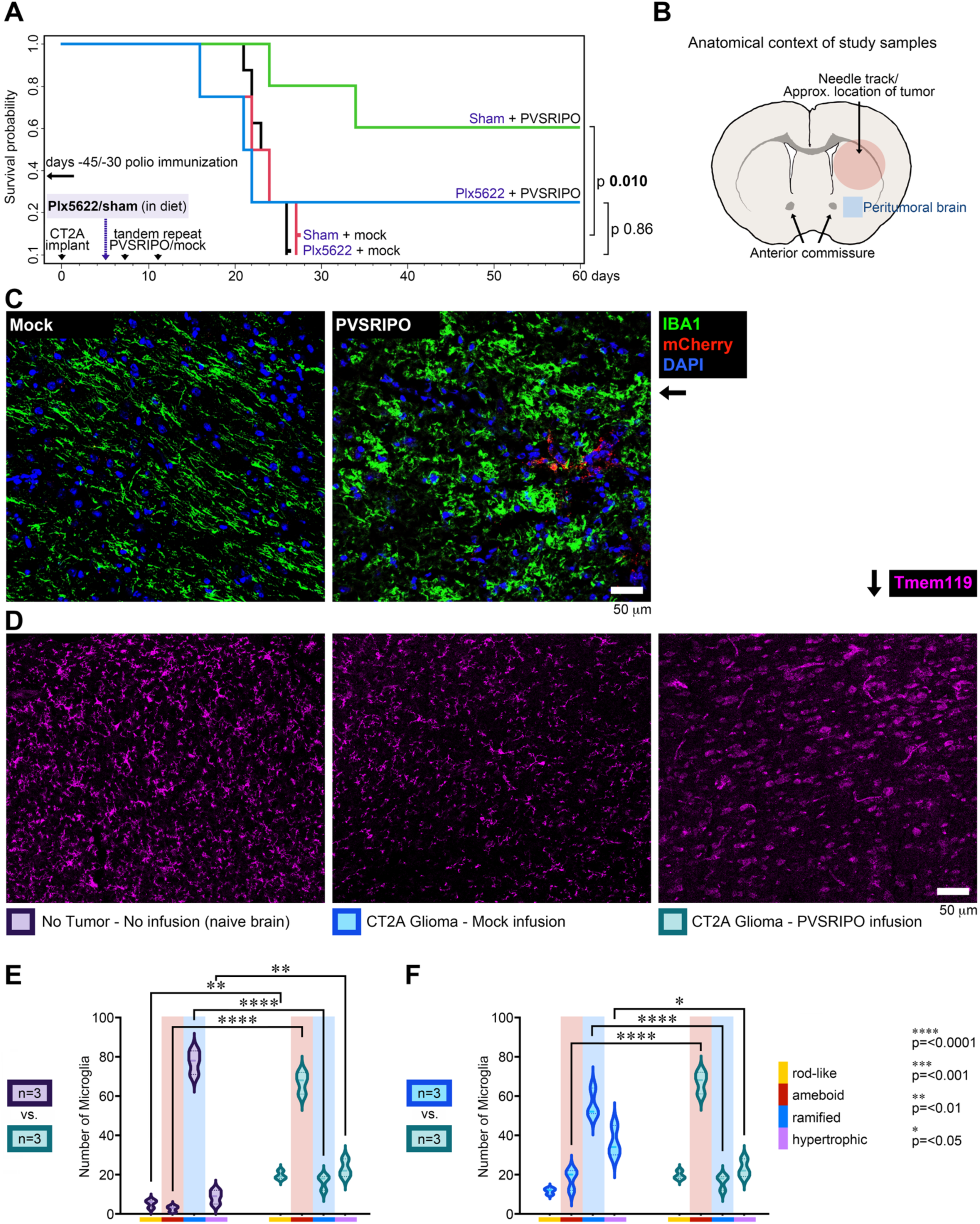
Disseminated microglia activation upon intracerebral PVSRIPO is essential for anti-glioma immunotherapy. **(A)** Survival after tandem-repeat intratumor PVSRIPO (5 x 10^7^ pfu) vs. mock (vehicle) in poliovirus-immunized *hCD155*-tg mice bearing CT2A^hCD155^ gliomas fed a Plx5622/sham-containing diet as shown. Survival curves were compared using a log-rank Mantel-Cox test. **(B)** Anatomical context of study samples. Stereotactic procedures for implanting tumor and inoculating virus used coordinates as reported previously (Yang et al., 2023) to target the central striatum. The blue (peritumoral brain) square indicates the approximate location of ipsilateral (uninjected) brain, distant from the needle track, examined throughout this work. **(C)** Validating the MicrogliaMorphology tool (see extended data in Figure S1). Iba1-stained peritumoral brain sections from CT2A^mCherry^-bearing mouse brains treated with mock vs. PVSRIPO (5 x 10^7^ pfu) were subjected to morphology analyses. Occasionally, mCherry^+^ tumor satellites were detected in peritumoral brain, as expected (eg. in the image shown in the right panel). **(D)** Tissue sections from naïve, non-infused *hCD155*-tg mouse brain (left), or from CT2A glioma-bearing *hCD155*-tg mice 48h after single stereotactic infusion of mock/PVSRIPO (5 x 10^7^ pfu; day 7 post tumor implantation) were stained with Tmem119 for morphology analyses. **(E, F)** Distribution of microglia shape prototypes comparing naïve to PVSRIPO-infused glioma-bearing brain (E) and mock-infused to PVSRIPO-infused glioma-bearing brain (F). Morphology data were analyzed by two-way ANOVA with Šídák’s multiple comparisons test.

To model malignant glioma we use C57Bl6 mice transgenic for <u>h</u>uman <u>p</u>oliovirus receptor [*PVR*; a.k.a., human (h)CD155]. *hCD155*-tg mice recapitulate hCD155 tissue-and cell type-specific distribution and develop classic paralytic polio upon wild type virus challenge (Koike et al., 1991). Murine bone marrow-derived macrophages/DCs and primary explant human monocyte-derived macrophages/ DCs display similar susceptibility phenotypes to PVSRIPO infection, and respond with similar innate signaling-and cytokine release patterns (Mosaheb et al., 2020). *hCD155*-tg mice were implanted with syngeneic CT2A mouse glioma expressing hCD155 [CT2A^hCD155^; mimicking virtually universal ectopic hCD155 expression in human malignant glioma (Chandramohan et al., 2017; Merrill et al., 2004)] as reported previously (Yang et al., 2023). A Plx5622/sham diet was initiated 5 days after and maintained until study day 40; mice were treated with tandem-repeat intratumor PVSRIPO/ mock and monitored for survival (**Fig. 1A**). To mimic the clinical scenario with pre-existing anti-poliovirus immunity due to routine infant vaccination and in recognizing key immunotherapy contributions of anti-polio immunity recall in the brain, mice were polio-immunized (Brown et al., 2023). Plx5622 did not affect the clinical course after tumor implantation compared to sham-fed mice (**Fig. 1A**). It did, however, abolish the significant (*p* 0.010; **Fig. 1A**) survival extension after PVSRIPO; in the presence of Plx5622, infusion of PVSRIPO lacked significant antitumor effects (*p* 0.86; **Fig. 1A**).

### PVSRIPO causes profound, disseminated changes in microglia morphology

The brunt of the microglia response identified in neuropathologist review of PVSRIPO-treated mouse glioma concerned peritumoral brain (Yang et al., 2023). Glioma myeloid infiltrates mount robust inflammatory reactions to PVSRIPO therapy (Yang et al., 2023), but these are difficult to decipher histologically due to the dense, chaotic architecture of myeloid infiltrates. Thus, our analyses focused on peritumoral brain: normal CNS tissue surrounding the tumor, which occasionally contained glioma satellites (**Fig. 1B, C**). Peripheral monocytic infiltration after PVSRIPO is limited to the needle track/tumor proper; it was absent from the areas chosen for our studies in peritumoral brain (Yang et al., 2023).

For an unbiased, broad-based test of microglia responsiveness to PVSRIPO, we conducted high-throughput analyses of shape characteristics of individual cells, generating high-dimensional datasets that were analyzed as described by Kim *et al*. (2024). Microglia exhibit profound intrinsic plasticity and occupy a fluid continuum of adaptive phenotypes (Hammond et al., 2019) where changing cell morphology signals reactive adaptation (Karperien et al., 2013). Iba1 immuno-fluorescence (IF), reliably marking microglia cell bodies and processes (Schwabenland et al., 2021) in peritumoral brain (**Fig. 1C**; for training/validation of the image analysis pipeline in Fiji and R; **Materials and methods**) revealed changed Iba1^+^ cell morphology after single intratumor PVSRIPO vs. mock (**Fig. 1C**). Iba1 also marks infiltrating monocytes and perivascular BaM in parenchyma; absent monocytic infiltration in peritumoral brain (Yang et al., 2023) and absent vessel-tracking stain, indicate Iba1^+^ cells in our assay to be microglia. Quantitative shape analysis results from the validation dataset are presented in Figure S1.

To systematically assess microglia morphology, tissue sections from the region corresponding to peritumoral brain (**Fig. 1B**) from naïve, non-infused, non-glioma-bearing brain; mock-infused glioma-bearing brain; and (single) PVSRIPO-injected glioma-bearing brain were tested with Tmem119 IF (**Fig. 1D**) for quantitative morphological analyses of microglia. The most pronounced effects identified concerned the ratios of morphologies at perceived opposite ends of a hypothetical polarization spectrum (Paolicelli et al., 2022): “ramified” morphology, typifying homeostatic states in the mature brain, vs. “ameboid” morphology, which embodies a broadly enhanced state of reactiveness (**Fig. 1D, E**). The ∼77:3 and ∼56:18 ramified:ameboid ratios in naïve and (mock-infused) glioma-bearing brains, respectively, significantly changed to a ∼16:67 ratio in PVSRIPO-infused glioma-bearing brain (*p* <0.0001; **Fig. 1E, F**). These findings are consistent with widespread, functional microglia reprogramming after intracranial PVSRIPO infusion.

### PVSRIPO mediates non-cytopathogenic, extended vRNA replication in microglia

PVSRIPO has a neuron-incompetent phenotype; a dsRNA-binding protein 76-anchored ribonucleoprotein complex intercepts all translation via the HRV2 IRES in neurons (Dobrikov et al., 2022; Merrill et al., 2006; Merrill and Gromeier, 2006; Neplioueva et al., 2010). Polioviruses do not infect astrocytes/oligodendroglia. Thus, intracerebral PVSRIPO in *hCD155*-tg mice [5 x 10^7^ plaque forming units (pfu)], non-human primates [up to 5 x 10^9^ pfu (Dobrikova et al., 2012)] and human subjects (Desjardins et al., 2018) did not elicit viral glial/neuropathic damage, canonical triggers of microglia reactions (Chhatbar and Prinz, 2021). This, and disseminated/non-focal microglia responses to PVSRIPO (**Fig. 1C, D**) suggested microglia reactiveness to be due to direct infection. Indeed, changes in microglia phenotype and antitumor effects of PVSRIPO therapy relied on hCD155 expression in non-malignant mouse host cells (Brown et al., 2021; Yang et al., 2023).

Since the main product of PVSRIPO infection is replicating vRNA (Dobrikov et al., 2023), we tested microglia infection by tracking (-) and (+)strand PVSRIPO RNA (**Fig. 2**). We used CD11b^+^-beads to isolate microglia (Bordt et al., 2020) from *hCD155*-tg mouse brains and from primary human (epilepsy surgery) brain tissue using the same approach and reagents (see **Materials and methods**). We routinely recovered ∼90-95% CD45^hi^ CD11b^+^ cells (**Fig. 2A**) that also were Cx3cx1^+^ and MERTK^+^ (**Fig. 2B**), and Tmem119^+^ (**Fig. 2C**), consistent with microglia lineage, from brain tissue of either species. Reverse transcription-quantitative PCR (RT-qPCR) of (+)strand vRNA using total RNA isolated from mouse (**Fig. 2D**, left panel) or human (**Fig. 2D**, middle panel) microglia infected *ex vivo* with PVSRIPO indicated active vRNA replication. RT-qPCR using total RNA recovered from CD11b^+^-bead isolated microglia harvested from *hCD155*-tg mice after intracerebral infusion of PVSRIPO provided initial evidence for vRNA replication in microglia *in vivo* (**Fig. 2D**, right panel). Detection of vRNA at similar levels in ipsilateral and contralateral brain hemispheres is consistent with altered microglia morphology throughout the brain (**Fig. 1**), and with prior evidence of brain-wide PVSRIPO dissemination after intracerebral inoculation (Ludwig et al., 2025).

**Figure 2.**
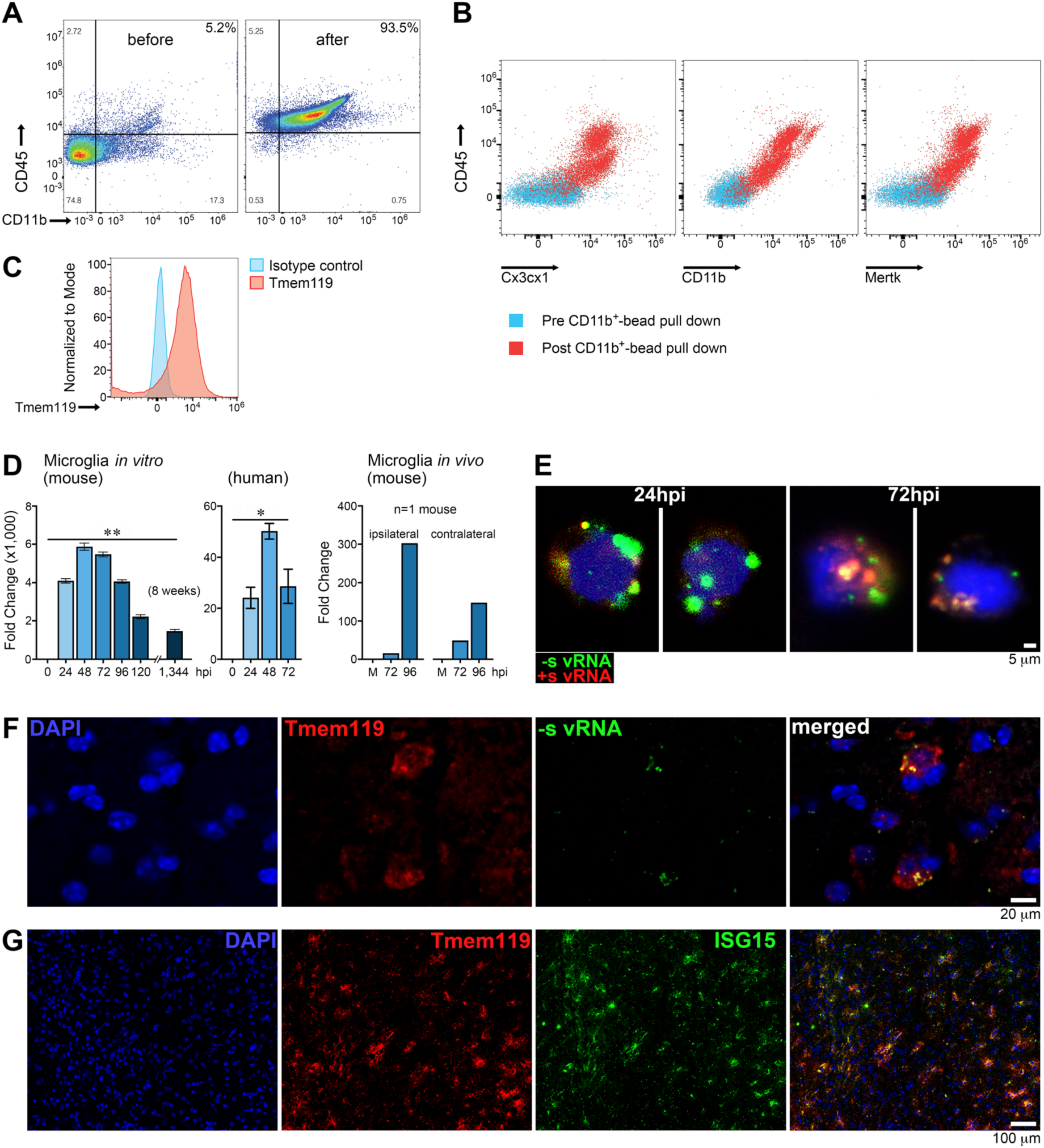
PVSRIPO directs infection of, non-cytopathogenic vRNA replication in, and IFN-I signaling in the microglial compartment *in vivo*. (A-C) Flow cytometry analyses of microglia before/after CD11b^+^-bead isolation from *hCD155*-tg mouse **(A, C)** or human brain tissue **(B)**. The same kit and identical procedures were used to process human/mouse brain tissue samples (see **Materials and methods**). Consistently, ∼90-95% of CD45^hi^, CD11b^+^, MERTK^+^, Cx3cx1^+^ **(A, B)** and Tmem119^+^ **(C)** cells were recovered from brain tissue of either species. **(D)** RT-qPCR analyses of total RNA from *hCD155*-tg murine or human microglia as shown. Microglia were either infected *ex vivo* [multiplicity of infection (MOI) 10; left and middle panels], or CD11b^+^-bead isolated after infection *in vivo* [single intracerebral inoculation of PVSRIPO (5 x 10^7^ pfu); right panel]. Ipsilateral (to the site of virus inoculation) and contralateral hemispheres were processed separately. vRNA levels in mouse microglia were analyzed by two-tailed Mann-Whitney test on ΔΔCt values at 8 weeks pi (n=5; left panel); the same test was used to analyze vRNA levels in human microglia at 72hpi (n=3; right middle panel). **(E)** HCR-FISH analyses of vRNA species in human CD11b^+^-bead isolated microglia infected *ex vivo* with PVSRIPO (MOI 10). Two individual microglia cells at 24 and 72hpi, each, are shown. **(F-G)** IF and HCR-FISH analyses in non-tumor-bearing *hCD155*-tg mouse brains 48h post single intracerebral PVSRIPO infusion (5 x 10^7^ pfu). Tmem119^+^ microglia stain positive for (-)strand vRNA **(F)** with widespread ISG15 induction in the microglial compartment **(G)**.

Hybridization chain reaction-fluorescence *in situ* hybridization (HCR-FISH) for microscopic imaging of vRNA in *ex vivo*-infected primary explant human microglia confirmed these observations (**Fig. 2E**): (-)strand vRNA (the initial vRNA replication product) in the characteristic distribution of viral replication organelles (Chen et al., 2015) at 24 hours post infection (hpi) gave rise to (+)strand vRNA (72hpi; **Fig. 2E**). Detection of (-)strand vRNA intermediates provided unambiguous evidence for PVSRIPO direct infection and active vRNA replication in human microglia. Consistent with prior findings in human monocyte-derived DCs (Brown et al., 2017), PVSRIPO infection of human/mouse microglia was non-cytopathogenic; indeed, microscopic analyses of microglia cultures showed enhanced adhesion/vitality in (+)strand vRNA+ microglia at eight weeks after infection (**Fig. 2D**) compared to mock-infected cultures.

### vRNA replication and disseminated IFN stimulated gene expression in the microglial compartment after PVSRIPO infusion *in vivo*

To document the PVSRIPO:microglia relationship *in vivo*, (non-tumor-bearing) *hCD155*-tg mice were treated with single intracerebral infusion of PVSRIPO using our standard stereotactic coordinates (**Fig. 1B**), followed by HCR-FISH and immunofluorescence (IF) analyses of the region corresponding to peritumoral brain (48hpi; **Fig. 2F, G**). Tmem119^+^ microglia co-stained for (-)strand vRNA with a signal distribution reminiscent of human microglia infected *ex vivo* (**Fig. 2F**, compare **Fig. 2E**), indicating active PVSRIPO RNA replication in microglia *in vivo*. This was accompanied by profuse IFN stimulated gene (ISG) 15 (ISG15) induction in a pattern almost entirely overlapping with Tmem119 signal (**Fig. 2G**).

### Single nuclei (sn)RNAseq of primary *ex vivo* human glioma tissue fragments

To gain insight into cellular responses to PVSRIPO in intact, heterogeneous, human CNS tissue in an unbiased global manner, we performed snRNAseq of human GBM patient *ex vivo* tissue slices treated with PVSRIPO vs. mock (**Figs. 3, 4**). We opted for this system because PVSRIPO is under clinical investigation for its capacity to generate glioma immune surveillance in the CNS (Desjardins et al., 2018); pivotal insight into PVSRIPO cell-type specificity and innate signatures stems from this model (Brown et al., 2021); and there is a mandate for tests of a poliovirus recombinant, derived of a consummate human pathogen, in authentic human models. Fresh malignant glioma tissue was obtained immediately after surgical resection, transferred to the laboratory, and processed for testing as previously reported [(Brown et al., 2021); **Materials and methods**]. Fragments were processed for snRNAseq 24hpi after addition of PVSRIPO/mock.

**Figure 3.**
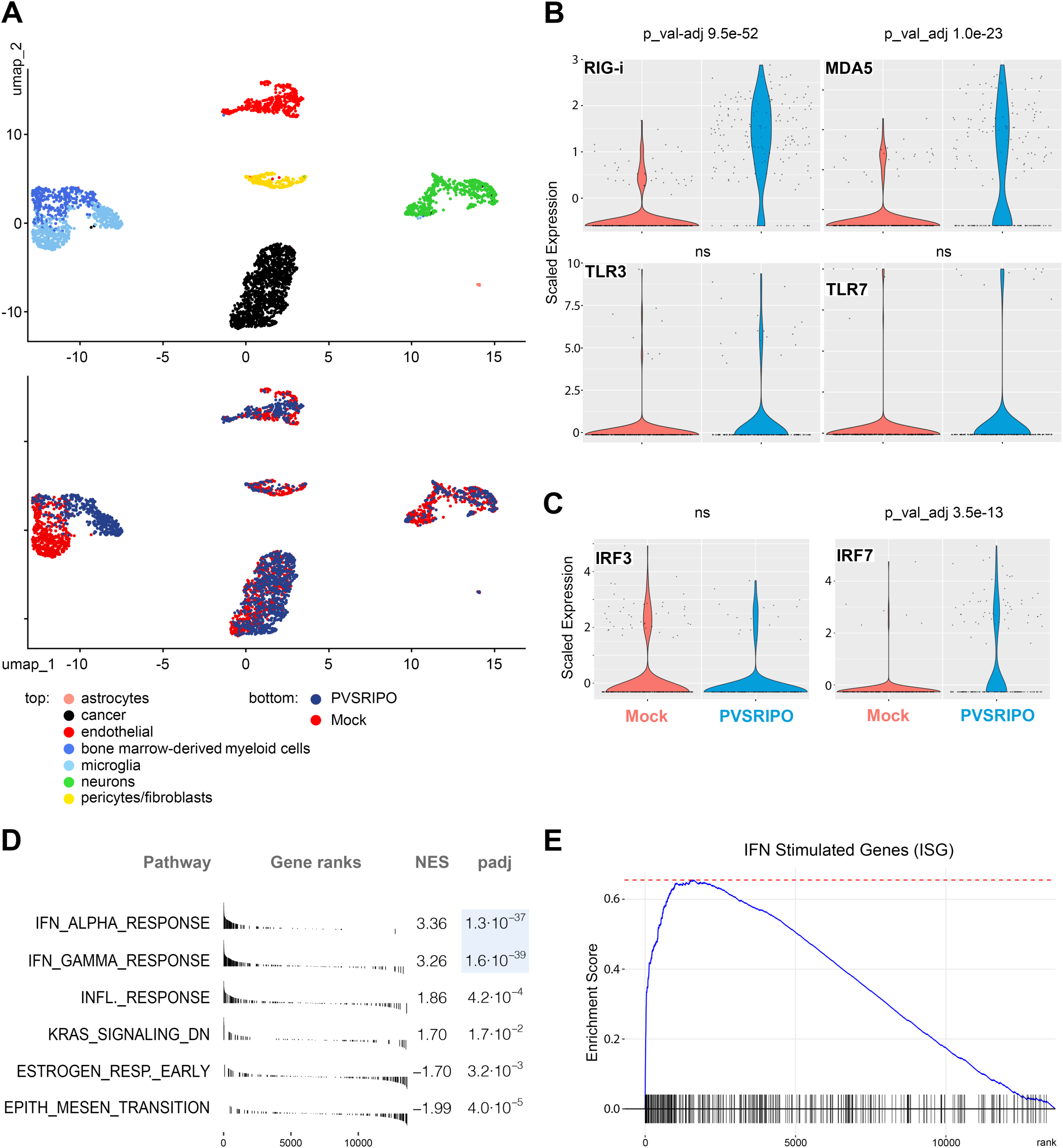
SnRNAseq identifies microglia as a principal site for the IFN-I/II dominant host response to PVSRIPO in the CNS. **(A)** UMAPs delineating cell type clusters (top) and an overlay of clustering treatment responses (PVSRIPO vs. mock; bottom) in GBM *ex vivo* slice cultures. See Table S1 for detailed gene lists informing cluster identification. **(B, C)** Induction of the cytoplasmic vRNA sensors RIG-I and MDA5 (B, top) vs. TLR3 and 7 (B, bottom) in microglia of mock vs. PVSRIPO-treated *ex vivo* slice cultures. Expression of IRF3 and IRF7 transcription factors in microglia of mock vs. PVSRIPO-treated *ex vivo* slice cultures (C). For statistical analyses, Bonferroni-adjusted *p* values from Wilcoxon rank-sum tests (Seurat `FindMarkers()`) were assessed. **(D, E)** GSEA hallmark analyses (D) and enrichment score for the canonical set of ISGs (E). Empirical *p* values were established by permutation testing (10,000 permutations) using a weighted Kolmogorov–Smirnov–like statistic (fgsea); adjusted using the Benjamini–Hochberg FDR method.

RNA expression of single nuclei derived from glioma *ex vivo* slices, normalized for treatment effect, revealed a large MPS cluster encompassing two smaller sub-clusters identified via Seurat. Gene set enrichment analysis (GSEA) using gene sets from the Molecular Signature Database (MSigDB; collection C8) was used to differentiate these sub-clusters as bone marrow-derived myeloid cells (bmdMC) and microglia (**Fig. 3A**; top; see also Figure 4 below). This MPS cluster demonstrated strong treatment-dependent separation when compared to other clusters, particularly in the microglial compartment (**Fig. 3A**; bottom).

**Figure 4.**
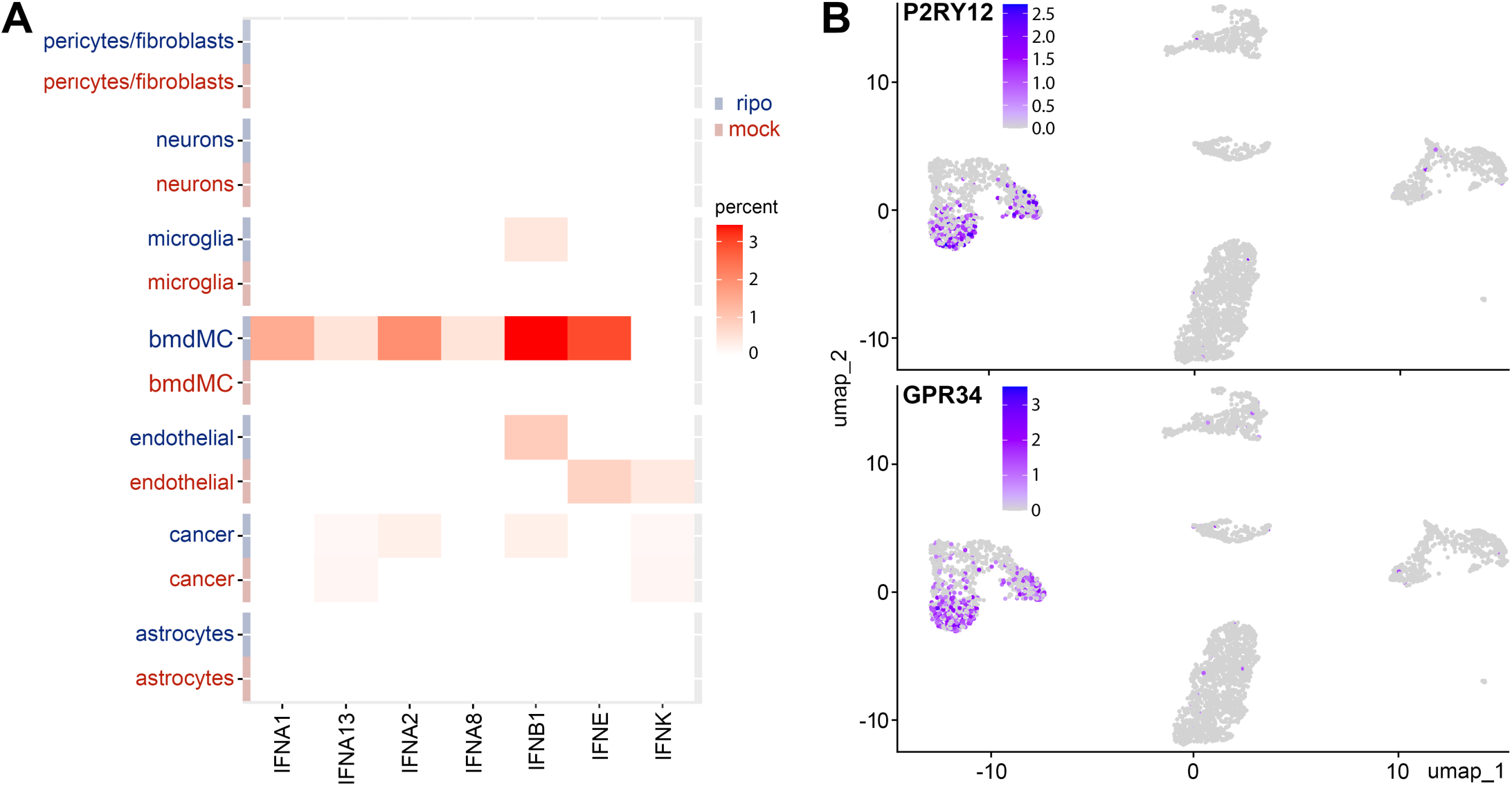
IFN-I release in GBM *ex vivo* tissue slice snRNAseq cell clusters. **(A)** IFN-I transcript frequency in distinct cell clusters. All detectable IFN-Is are shown. **(B)** Differentiation of microglia and bmdMC clusters based on P2RY12 and GPR34 transcript frequency. See Table S1 for detailed gene lists informing microglia vs. bmdMC differentiation.

In PVSRIPO-treated microglia, the cytoplasmic vRNA sensors (and ISGs) RIG-I and MDA5 were significantly upregulated (*p* 9.5 x 10^-52^ and *p* 1.0 x 10^-23^, respectively; **Fig. 3B**, top) while the endolysosomal TLR3 and TLR7, which are not involved in PVSRIPO vRNA sensing/are not ISGs, were not (**Fig. 3B**, bottom). The hallmark IFN signaling response to PVSRIPO in mononuclear phagocytes is sustained IRF7 induction [corresponding with sustained IFNα release in human monocyte-derived macrophages; (Dobrikov et al., 2023)]; accordingly, IRF7 (ISG expressed at very low baseline levels in macrophages/microglia/myeloid DCs) was significantly induced in microglia in PVSRIPO-treated *ex vivo* slices (*p* 3.5 x 10^-13^; **Fig. 3C**). Meanwhile, levels of IRF3, expressed constitutively in myeloid cells, did not change upon PVSRIPO treatment (**Fig. 3C**).

GSEA using Hallmark pathway gene sets revealed dominant and significant enrichment of both IFNα and IFNγ response pathways in PVSRIPO-treated microglia (*p* 1.3 x 10^-37^ and *p* 1.6 x 10^-39^, respectively; **Fig. 3D**). This coincided with PVSRIPO-induced significant enrichment of canonical ISGs, reflecting robust and select activation of IFN-I and-II signaling pathways in these cells (**Fig. 3E**). This agrees with our HCR-FISH, RT-qPCR and imaging analyses of human and murine microglia *in vitro* and *in vivo*, showing non-cytopathogenic, extended vRNA replication in mononuclear phagocytes as the principal feature of the PVSRIPO:host relationship in the brain.

### PVSRIPO microglia transcriptional reprogramming lacks cytokine release

PVSRIPO-mediated IRF7 induction in microglia, implying IRF3 activation, may suggest IFN-I release as a result [IRF3/IRF7 are the master transcriptional regulators of *IFNA/IFNB* (Sato et al., 2000)]. We tested this by mapping detectable IFN-I transcripts to specific cell clusters in our *ex vivo* tissue slice snRNAseq data set (**Fig. 4**). Consistent with PVSRIPO’s cell type specificity, IFN-I transcripts were not detected in neuron/astrocyte clusters (**Fig. 4A**). We separated microglia/ bmdMC clusters based on microglia-focused expression, eg. of purinergic receptor P2RY12 (Sasaki et al., 2003) and probable G protein-coupled receptor (GPR34) (Bennett et al., 2016) (**Fig. 4B**). PVSRIPO’s hallmark innate signature, sustained IFNα release (Brown et al., 2021), was triggered in bmdMCs, but not microglia (**Fig. 4A**). IFNβ production, an initial IFN-I response to PVSRIPO infection that precedes IFNα release (Dobrikov et al., 2023), was observed (faintly) in microglia, as well as (strongly) in bmdMCs, endothelial cells and in the neoplastic compartment. Our findings indicate that transcriptional reprogramming of microglia upon PVSRIPO infection, evident as an IFN-I/II dominant GSEA signature (**Fig. 3**), drives internal ISG expression programs, but may not produce IFN-I release at levels observed in bmdMCs.

### PVSRIPO infection induces striking microglia phagocytic activity *in vitro* and *in vivo*

In line with their status as the “professional” phagocytes of CNS parenchyma, microglia roles in neurodevelopment, CNS homeostasis, and immune surveillance rely on phagocytic activity. Thus, it was pivotal to ascertain microglia phagocytosis activity upon innate reprogramming with PVSRIPO infection/vRNA replication. First, we assessed phagocytosis of (mouse) CT2A glioma cells by CD11b^+^-bead isolated, *hCD155*-tg mouse microglia after mock vs. PVSRIPO infection over time. CT2A cells were Cell Trace (CT)-labeled, UV-inactivated and “fed” to microglia at the same time of infection with mock/PVSRIPO, followed by flow cytometry analyses of CT^+^ microglia (**Fig. 5A**). We observed robust phagocytic activity, at levels ∼27-125-fold over mock, at 24-72hpi (**Fig. 5A**). Evidence for CT^+^ cell phagocytic uptake and break-up was visible in photomicrographs of PVSRIPO-infected microglia (**Fig. 5B**).

**Figure 5.**
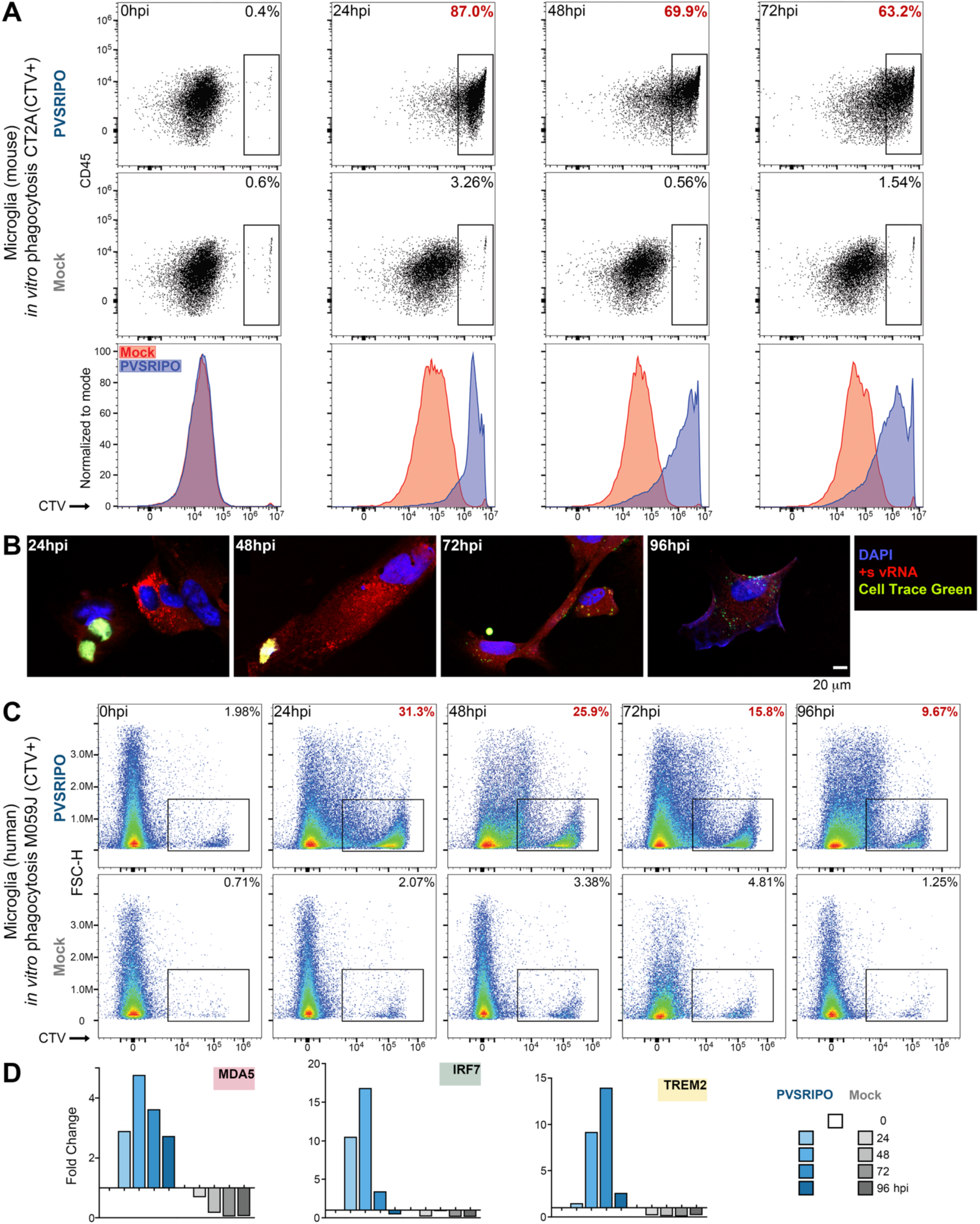
PVSRIPO infection induces robust phagocytic activation of primary explant human and murine microglia. **(A)** *In vitro* phagocytosis assay of CD11b^+^-bead isolated *hCD155*-tg mouse microglia. UV-inactivated, CT^+^ CT2A glioma cells were fed to microglia at the time of mock/ PVSRIPO (MOI 10) infection. Proportions of phagocytically active (CT^+^) microglia were recorded by flow cytometry (top rows); histograms compare PVSRIPO vs. mock treatment in single plots (bottom row). **(B)** Photomicrographs of CD11b^+^-bead isolated *hCD155*-tg mouse microglia, infected with PVSRIPO, engulfing and degrading CT^+^ CT2A glioma cells. **(C)** *In vitro* phagocytosis assay using CD11b^+^-bead isolated human microglia exposed to UV-inactivated, CT^+^ human M059J glioma cells using procedures as shown in panel A. Proportions of phagocytically active (CT^+^) microglia were recorded by flow cytometry. **(D)** Human microglia samples profiled for phagocytic activity (C) were tested for induction of MDA5, IRF7 (see Figure 3) and TREM2 by RT-qPCR.

A similar outcome was observed with primary explant human microglia (from epilepsy surgery brain) exposed to human CT^+^ M059J glioma cells (**Fig. 5C**). Phagocytic activity peaked at 24hpi with ∼31% CT^+^ microglia in the culture, a ∼15-fold increase over mock (**Fig. 5C**). PVSRIPO-mediated human microglia phagocytic activity was accompanied by induction of MDA5, IRF7, and the Triggering Receptor Expressed on Myeloid cells 2 (TREM2; **Fig. 5D**), implicated in regulating phagocytic activity in microglia (Takahashi et al., 2005).

Similar levels of phagocytic activity were observed *in vivo*, upon PVSRIPO infusion into CT2A^mCherry^-bearing *hCD155*-tg mouse brains (**Fig. 6A**). Flow cytometry of CD11b^+^-bead isolated microglia, recovered 48h post single PVSRIPO/mock treatment, revealed that ∼47% of recovered microglia were mCherry^+^, vs. ∼7% in mock-treated animals. This coincided with induction of MHC II, MERTK and CD86 in microglia from PVSRIPO vs. mock-treated brains (**Fig. 6A-C**). mCherry tracer engulfment by microglia was also evident by co-localizing IF of Tmem119 *in vivo* (**Fig. 6D**).

**Figure 6.**
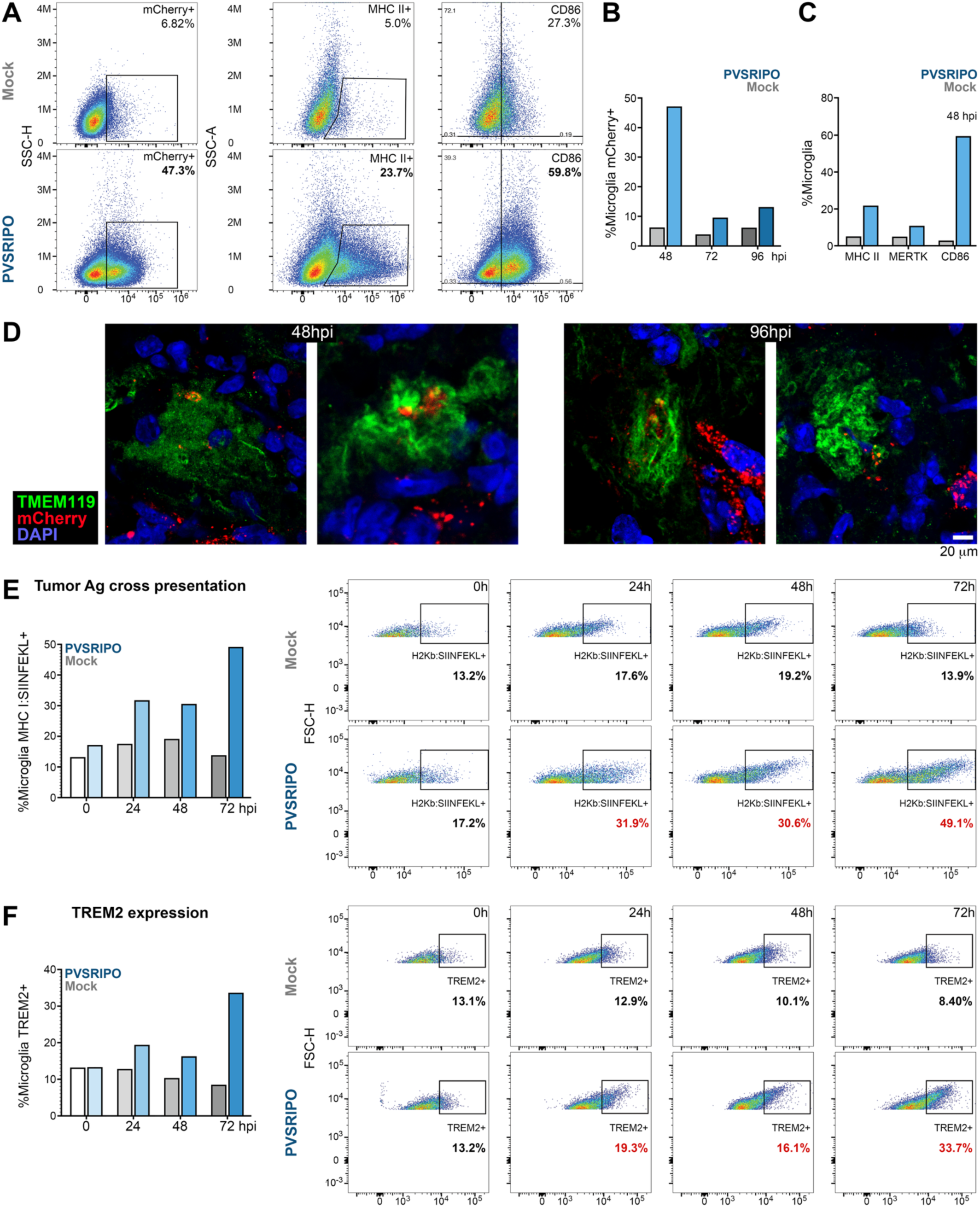
PVSRIPO infection-triggered phagocytic activity *in vitro* and *in vivo* leads to cross presentation of ingested antigens. (A-C) *hCD155*-tg mice bearing orthotopic CT2A^hCD155^ malignant gliomas, labeled with mCherry tracer, were treated with a single stereotactic intratumor PVSRIPO infusion (5 x 10^7^ pfu). Mice were euthanized at 48h, 72h or 96h post treatment, the tumor-bearing hemisphere was dissected and processed for CD11b^+^-bead microglia isolation for flow cytometric assessment of phagocytic activity and maturation marker expression. At the peak 48hpi interval, phagocytic activity was evident as ∼47% of microglia staining positive for mCherry (vs. ∼6.8% in the mock-treated control), associated with elevated frequencies of MHC II^+^, MERTK^+^, and/or CD86^+^ microglia (A-C). Phagocytic activity was also detected by imaging of (peritumoral) brain sections, with Tmem119^+^ microglia engulfing the mCherry tracer at 48 and 96hpi **(D)**. **(E)** Tumor antigen cross presentation by PVSRIPO-infected microglia was assessed by methods analogous to assays shown in Figure 5: UV-inactivated B16^OVA^ cells were fed to CD11b^+^-bead-isolated *hCD155*-tg mouse microglia simultaneously infected with PVSRIPO (MOI 10) vs. mock. Flow cytometry assays with an antibody specific for the H2Kb:SIINFEKL complex revealed proportions of cross presenting microglia amongst all microglia in the assay. **(F)** In parallel to cross presentation, the frequency of TREM2^+^ microglia was evaluated in the samples tested in panel E.

### PVSRIPO-infected microglia phagocytose and cross-present tumor antigen at levels exceeding inflammatory mediators

PVSRIPO infection of MPS cells induces: the immunoproteasome, peptide transport machinery [eg. transporter-associated protein 1 (TAP1) (Brown et al., 2017)], MHC I and the lymph node homing receptor, CCR7 (Mosaheb et al., 2020). We therefore tested if tumor cell phagocytosis by PVSRIPO-infected microglia triggers cross presentation of tumor antigens. The murine microglia phagocytosis assay in Figure 5A was performed with CT-labeled, UV-inactivated B16 melanoma cells expressing OVA. Microglia were mock/PVSRIPO-infected, exposed to B16^OVA^ bait, and tested by flow cytometry with an antibody specific for H2Kb:SIINFEKL complexes (**Fig. 6E**). Robust (OVA MHC class I-restricted epitope) SIINFEKL cross presentation in PVSRIPO-infected microglia (**Fig. 6E**) was accompanied by TREM2 induction (**Fig. 6F**), which was also observed by RT-qPCR in human microglia after PVSRIPO infection (**Fig. 5D**). TREM2, a sensor of damage-associated molecular patterns (eg. apoptotic cells, phospholipids, apolipoprotein E, Aβ) (Wang et al., 2022), is expressed in a pattern matching CD155 (microglia, macrophages, myeloid DCs), and has roles in modulating phagocytic activity, innate inflammatory states and metabolism in the MPS.

We assessed PVSRIPO-triggered B16^OVA^ phagocytosis and SIINFEKL cross presentation in murine microglia *in vitro*, side-by-side with other, functionally relevant comparators: intracellular high molecular weight poly(I:C) and recombinant IFNγ. Transfected poly(I:C) (LyoVec complex; **Materials and methods**), like PVSRIPO RNA, is sensed by MDA5 (Gitlin et al., 2006), in addition to TLR3 (Alexopoulou et al., 2001). IFNγ is a master regulator of the immunoproteasome, peptide loading complex and cross presentation in MPS cells (Schroder et al., 2004). The assay was conducted as described for Figure 5E; mock/poly(I:C)/IFNγ/PVSRIPO treatment was initiated at the same time of adding bait UV-inactivated B16^OVA^ cells to the cultures. PVSRIPO-infected microglia exhibited significantly elevated phagocytic activity (eg. compared to mock at 48hpi *p* <0.001, **Fig. 7A, C**) and SIINFEKL cross-presentation (*ibid*. **Fig. 7B, C**) compared to mock/poly(I:C)/IFNγ at all intervals examined.

**Figure 7.**
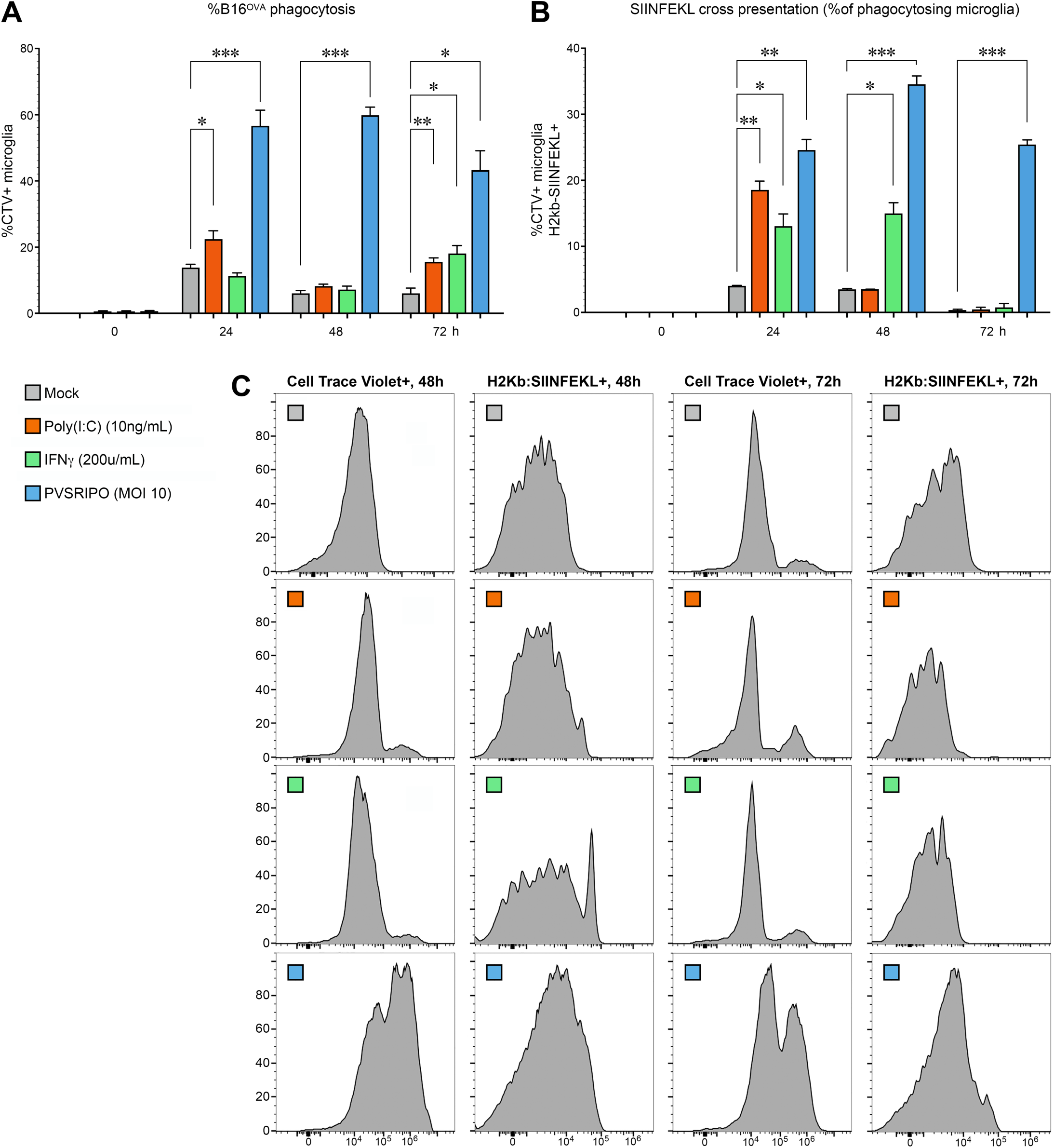
**Microglia phagocytic activity and antigen cross presentation triggered by treatment with mock, poly(I:C), recombinant IFN**γ**, and PVSRIPO.** The phagocytosis/cross presentation assay template with microglia exposure to B16^OVA^, described in Figure 6E, was repeated with juxtaposing PVSRIPO to mock, (transfected) poly(I:C), and recombinant IFNγ. Phagocytosis of CD11b^+^-bead-isolated, *hCD155*-tg mouse microglia was determined by flow cytometry analyses of CT-label **(A**, **C)**; antigen cross presentation was tested by flow cytometry for H2Kb:SIINFEKL complexes **(B**, **C)**; cells were tested 0h, 24h, 48h and 72h post treatment; flow cytometry histograms for phagocytosis/cross presentation at 48h and 72h post treatment are shown **(C)**. **(A**, **B)** Data were analyzed by Shapiro-Wilk test and 2-way ANOVA with multiple comparisons.

### PVSRIPO-triggered microglia reprogramming is associated with sweeping amyloid-beta clearance in an injectable amyloid oligomer model

Our data imply a marked capacity of PVSRIPO RNA replication to engage microglia in phagocytic uptake of neoplastic material. We therefore examined PVSRIPO-directed phagocytosis of burdensome debris in CNS parenchyma of another clinically relevant entity: Aβ deposits associated with Alzheimer’s Disease (**Figs. 8, 9**). Implantation of (*in vitro* preaggregated) Aβ(1-42) HiLyte Alexa Fluor (AF)488 oligomers (Aβ HiLyte) yielded Aβ deposits by day 28 (**Fig. 8A**), consistent with prior reports (Baleriola et al., 2014; Brouillette et al., 2012). At this interval, Aβ HiLyte-bearing mice received single stereotactic inoculation of PVSRIPO (5 x 10^7^ pfu) vs. mock (vehicle) and were euthanized 48h thereafter. Microglia were CD11b^+^-bead isolated from four PVSRIPO or mock-treated mice, respectively, and assessed by flow cytometry for intracellular AF488 label. This revealed significantly enhanced microglial Aβ uptake (*p* <0.05, **Fig. 8B**); induction of CD40, CD86 and CD68 in microglia (**Fig. 8C**); and enhanced microglia MHC II expression (**Fig. 8D**) in the PVSRIPO cohort vs. mock-treated controls. Further analyses of CD11b^+^-bead isolated microglia recovered from Aβ HiLyte-injected animals at day 70, 48h post single PVSRIPO/mock therapy, provided additional insight into the breadth and depth of microglia reprogramming (**Fig. 8E**). Our analyses revealed three microglia clusters, CD45^lo^, ^-med^, and ^-hi^ in PVSRIPO-treated mouse brains; the CD45^hi^ cluster was all but absent in mock-treated animals (**Fig. 8E**). Flow cytometry analyses revealed significant induction of ApoE, CD86, Ki67, and MERTK in microglia of the CD45^lo^ and ^-med^ clusters from PVSRIPO vs. mock-treated mice (*p* <0.05; **Fig. 8F**). The CD45^hi^ cluster from PVSRIPO-treated animals stood out with >80% Ki-67^+^ positive cells, indicating a sizeable population of proliferating microglia in PVSRIPO-treated mouse brains, consistent with prior reported results (Yang et al., 2023). All microglia clusters isolated from PVSRIPO-treated animals exhibited significantly elevated phagocytic activity against Aβ HiLyte (*p* <0.05; **Fig. 8G**), compared to mock-treated animals. As indicated by microglia morphology analyses (**Fig. 1**), by the dissemination of vRNA replication and ISG induction (**Fig. 2F, G**), snRNAseq results (**Fig. 3**), and by *in vivo* phagocytosis results (**Fig. 6A**), these findings indicate widespread engagement of the microglial compartment upon PVSRIPO immunotherapy *in vivo*.

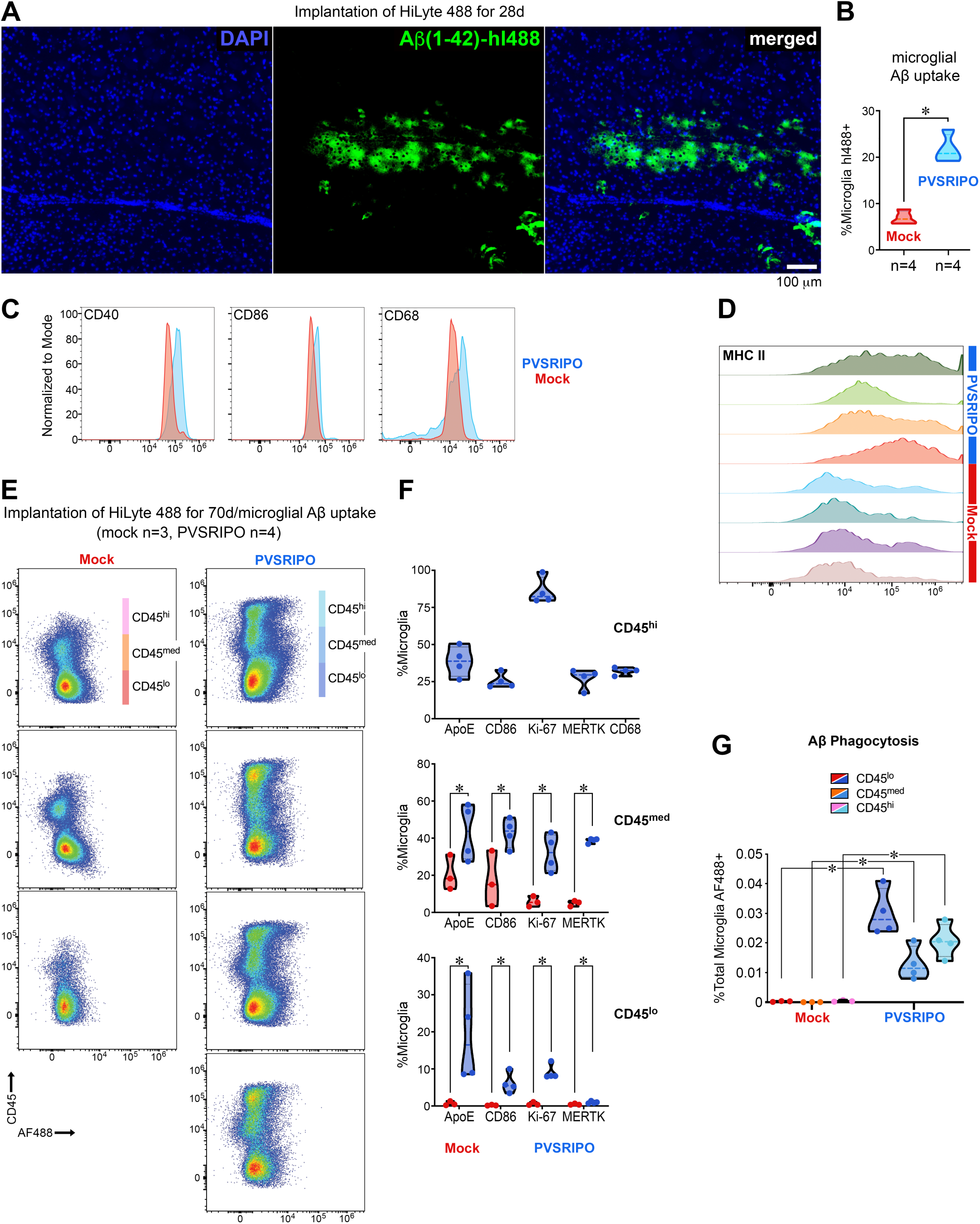
injectable Aβ(1-42) HiLyte AF488 oligomer accumulation model in the brain. **(A)** Aβ HiLyte (pre-incubated at 37°C, 72h) was stereotactically implanted in *hCD155*-tg mice (2.5μg in 5μl) using coordinates shown in Figure 1B. The photomicrographs depict deposits 28 days after Aβ implantation at the site of injection. **(B)** In a separate assay, Aβ HiLyte-implanted mice (28 days) received single stereotacting injections of mock (n=4) or PVSRIPO (5 x 10^7^ pfu; n=4). Brains were dissected 48h after virus administration, microglia were recovered by CD11b^+^-bead isolation and tested for AF488 uptake by flow cytometry. Aggregate results indicate significantly elevated Aβ phagocytosis by microglia *in vivo* (*p* <0.05). Data were analyzed by two-tailed Mann-Whitney test. **(C, D)** Microglia *in vivo* Aβ phagocytosis was accompanied by CD40, CD86 and CD68 induction (C) and MHC II upregulation *in vivo* (D). **(E)** In another separate assay, Aβ HiLyte-implanted mice (70 days) were treated (mock, n=3; PVSRIPO, n=4) and microglia isolated as in (B-D) for deeper flow cytometry-based phenotypic analyses. Gating for CD45 and AF488 revealed three clusters with distinct CD45 expression (CD45^lo/med/hi^) in PVSRIPO-treated brains; the CD45^hi^ cluster was virtually absent from mock-treated animals. **(F)** Flow cytometry analyses of microglia immune surveillance markers ApoE, CD86, Ki67, MERTK and CD68 in the CD45^lo/med/hi^ clusters. Significant elevation of all markers tested in the CD45^med/lo^ clusters occurred in the PVSRIPO compared to the mock cohort (*p* <0.05); the CD45^hi^ cluster was only analyzed in the PVSRIPO cohort. Data were analyzed by Shapiro-Wilk test and multiple unpaired t-test. **(G)** Flow cytometry analyses of AF488 phagocytosis in the distinct CD45^hi/med/lo^ clusters. Phagocytic activity was significantly elevated in all clusters of the PVSRIPO cohort (*p* <0.05).

We next investigated the effect of PVSRIPO (vs. mock) on Aβ HiLyte burden at the CNS-wide level and in the cervical (brain draining) lymph nodes (cLN; **Fig. 9**). Mice received Aβ HiLyte injections as described for Figure 8, were maintained for ten weeks thereafter, followed by single intracerebral inoculation of mock (**Fig. 9A**) vs. PVSRIPO (**Fig. 9B**) using the same stereotactic coordinates. Brains from one mouse in each cohort were dissected 1 and 3 weeks post mock/PVSRIPO therapy and stained with A11 antibody (recognizing amyloid oligomers), since spontaneous AF488 fluorescence of Aβ HiLyte was not visible at lower magnification for display of whole-brain images. The brains recovered from PVSRIPO-treated mice 7 days after virus/mock inoculation (70 days post Aβ HiLyte injection plus 7 days post PVSRIPO/mock treatment) showed substantial relief from parenchymal Aβ deposits compared to the heavily burdened brain of mock-treated animals (**Fig. 9A, B**). Brains dissected 14 days later (70 days post Aβ HiLyte injection plus 21 days post PVSRIPO/mock treatment) revealed a similar clearance phenomenon, indicating durable Aβ HiLyte clearance after PVSRIPO immunotherapy (**Fig. 9C, D**). Dissection of deep cLN, ipsilateral to Aβ HiLyte/PVSRIPO/mock stereotactic inoculation, revealed *ex*-CNS trafficking of Aβ HiLyte with meningeal lymphatic drainage (**Fig. 9E**), indicating the possibility of PVSRIPO-induced meningeal lymphatic uptake as a means of Aβ clearance from the brain.

**Figure 9.**
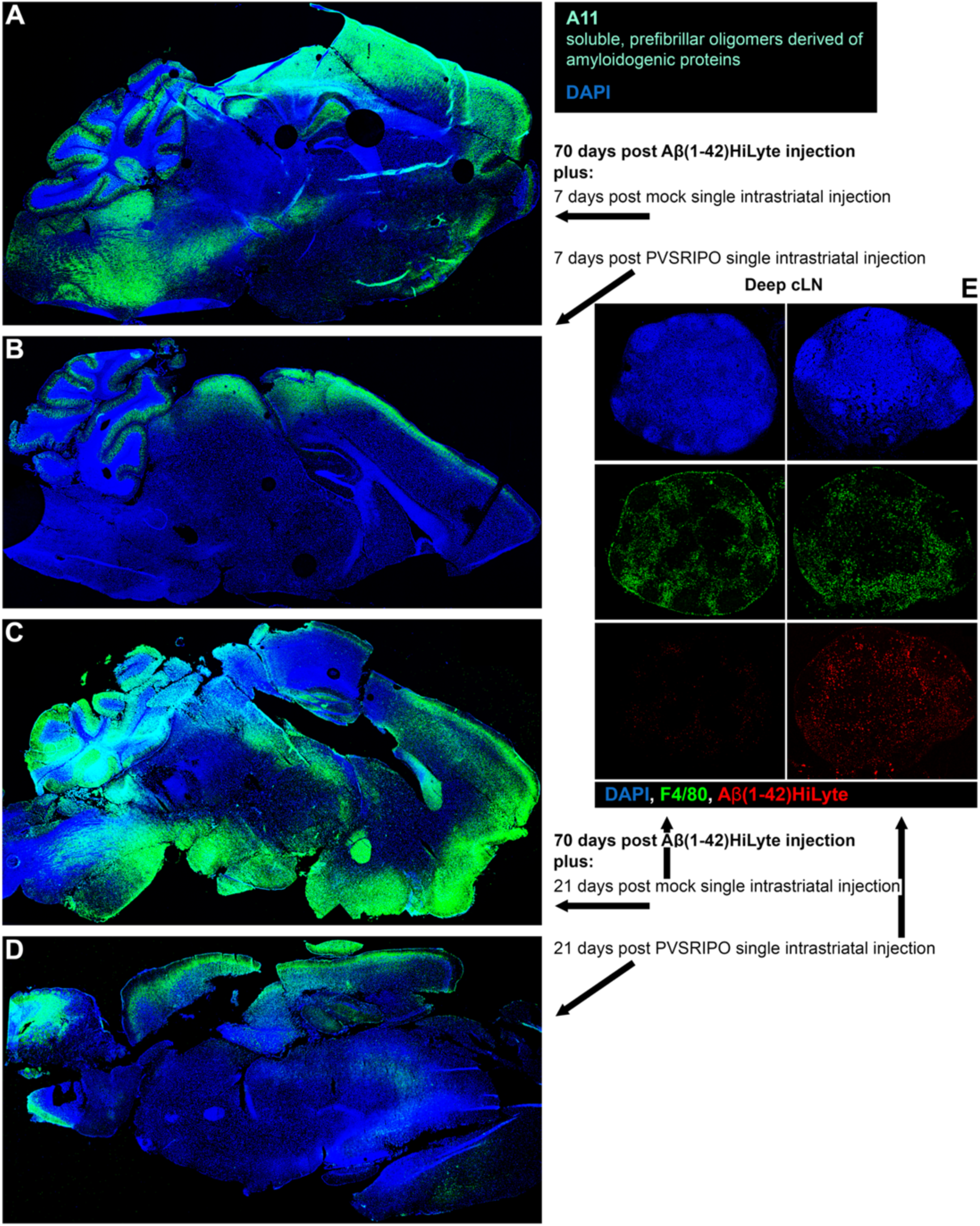
PVSRIPO clears parenchymal. **A**β **deposits and induces meningeal lymphatic trafficking of A**β **in an injectable oligomer model. (A-D)** Mice received stereotactic intracerebral implants of Aβ HiLyte (2.5μg in 5μl) using the coordinates shown in Figure 1B. Implanted mice were maintained for 70 days before single stereotactic inoculation of mock (A) or PVSRIPO (5 x 10^7^ pfu) (B) using the same coordinates. Brains of mice were harvested 7 days (A, B) or 21 days (C, D) after PVSRIPO/mock treatment and sagittal cryosections (5μm) were processed for A11 staining. **(E)** Deep cLN from the same animals shown in panels C (mock-treated) and D (PVSRIPO-treated) were stained as shown; spontaneous AF488 signal of Aβ HiLyte was imaged.

## Discussion

Microglia function has been described mainly as reactive to external cues, such as cell damage and danger-, mechanosensory-or inflammatory mediator-triggered adaptation. Here we show viral RNA replication-mediated microglia reprogramming that induces phagocytic, antigen cross presentation, and parenchymal debris clearance activities. MPS cells coordinate immune responses by sensing-and integrating a multitude of specific, disparate signals. Cell external stimuli are likely insufficient to convey the full breadth of such context; indeed, *endogenous* PRR activation/innate signaling was shown to be necessary for fully engaging DC function (Sporri and Reis e Sousa, 2005).

We discovered MDA5-triggered, IRF3/IRF7 dominant, select IFN-I/II transcriptional reprogramming marked by a robust ISG signature in PVSRIPO-infected human microglia, acompanied by highly restricted IFN-I secretion. PVSRIPO’s unique, “polar” IRF3/IRF7 innate footprint may be a result of selective MDA5 agonism, TBK1:IKKε dominant signaling (Brown et al., 2021), the lack of viral host innate interference, and lacking cytotoxicity [reviewed in (Gromeier and Brown, 2025)]. IRF3 activation can directly induce ISGs, bypassing IFN-I release and paracrine IFNAR signaling (Grandvaux et al., 2002). Of the five “classic” IFN-I generating PRRs, by virtue of their ability to activate IRF3/IRF7 (Liu et al., 2015), three (MDA5, RIG-I, TLR3) are dedicated to vRNA sensing. Thus, natural RNA virus infection may be particularly apt to deliver innate stimuli with spatiotemporal context and quality capable of enabling MPS immune surveillance (Freigang et al., 2005). This reflects the quintessential foreign and danger signatures of replicating vRNA (Matzinger, 2002), eg. of the lethal encephalitogenic RNA viruses. Thus, an internal innate reprogramming phenotype induced by PVSRIPO, with “IFN-I polarity” devoid of proinflammatory NFκB cytokine signatures (Brown et al., 2021), may protect bystander brain tissue from toxic neuroinflammation while delivering the immune-stimulatory profile of the IFN-I program to the microglial compartment. Indeed, neurotoxic microglial inflammation was linked specifically to NFκB cytokine release (TNF, IL-1β, IL-10, IL-17) upon TLR sensing of Aβ deposits (Jin et al., 2008). Our findings in microglia reverberate with clinical observations of absent neuroinflammatory toxicity with intracerebral PVSRIPO administration in patients (Desjardins et al., 2018; Thompson et al., 2023), in non-human primates (Dobrikova et al., 2012) or in *hCD155*-tg mice treated in this study and previously (Ludwig et al., 2025; Yang et al., 2023).

Polioviruses naturally target macrophages (both tissue-resident and monocyte-derived) and myeloid DCs, which exhibit the highest expression of the poliovirus receptor CD155 among immune cells (Monaco et al., 2019), at their portal of entry (Shen et al., 2017). This tropism may be decisive for poliovirus’ host immune interference strategy, as it may enable homing to the lymphatic system (Ludwig et al., 2025; Shen et al., 2017), the main virus replication reservoir (Bodian, 1955; Sabin, 1956) and cytolytic targeting of DCs and high endothelial venules in lymph nodes, the site where adaptive antiviral immunity is primed (Ludwig et al., 2025). PVSRIPO retains poliovirus’ cell type-specificity and capacity for installing vRNA replication organelles in infected macrophages/DCs (in the presence of robust innate antiviral IFN-I responses), but it lacks the ability to overcome host defenses, cause cytopathogenic damage, or direct virus proliferation.

Prior studies discovered decisive roles for vRNA replication-triggered MDA5 signaling in macrophages/DCs in directing CD4^+^T cell T_H_1 polarization and CD8^+^T cell priming (De Giovanni et al., 2020; Wang et al., 2012), emphasizing the pivotal role of spatiotemporal context of innate signaling in determining immune function in MPS compartments (Trinchieri, 2010). PVSRIPO’s MDA5 agonism and IRF3/IRF7 polar signature are likely part of an ancient protective antiviral program, driven by poliovirus:human co-evolution over millenia. In contrast, aberrant MDA5 (or cGAS-STING) activation with uncontrolled IFN-I release in the CNS, observed with Aicardi-Goutieres Syndrome-associated disruption of nucleic adic metabolism, provides a very different, pathogenic context with highly neurotoxic outcomes (Liu and Ying, 2023).

Our investigations revealed robust, sustained phagocytic activity *in vitro* and *in vivo*, antigen cross presentation and parenchymal debris clearance *in vivo* induced by PVSRIPO infection. The magnitude and duration of PVSRIPO’s effects on these immune surveillance functions was notable and exceeded what was seen *in vitro* with the TLR3/MDA5 agonist (transfected) poly(I:C) or a “master” inducer of antigen cross presentation, IFNγ. We attribute this differential to PVSRIPO’s non-cytopathogenic profile with active vRNA replication within microglia, and the quality and depth of innate stimulation this entails.

Poliovirus (capsid diameter ∼27nm) is naturally highly CNS-invasive; yet, the range of PVSRIPO dissemination within the CNS is remarkable, given that the virus is non-cytopathogenic and incapable of driving virus proliferation in the CNS (Ludwig et al., 2025). Non-lytic dissemination via exosomal vehicles, described for wild-type poliovirus (Chen et al., 2015), also occurs with PVSRIPO *in vitro* (Ludwig et al., 2025). We were unable to demonstrate non-lytic cell-to-cell spread in a relevant *in vivo* context, due to technical limitations in recovery, purification and experimental use of exosomal material. Yet, such modes of dissemination may explain PVSRIPO’s reach in the brain.

Our findings of oligomeric neurotoxic debris clearance in CNS parenchyma and Aβ meningeal lymphatic drainage in the injectable Aβ HiLyte model *in vivo* raise intriguing potential for PVSRIPO in the immunotherapy of neurodegenerative disease. It is highly likely that the meningeal lymphatic system (Louveau et al., 2015), which is prominentely targeted by the naturally lymphotropic PVSRIPO (Ludwig et al., 2025), participates in the CNS immunotherapy response to PVSRIPO. A mechanistic link of meningeal lymphatic drainage and microglia responsiveness to misfolded protein aggregation has been demonstrated previously (Da Mesquita et al., 2018; Da Mesquita et al., 2021). Indeed, the meningeal lymphatic system may broadly control immune surveillance in the CNS (Louveau et al., 2018). Deciphering the involvement of the CNS-resident phagocytic system at large in PVSRIPO immunotherapy, and unraveling microglia function within a more relevant pathobiological context will require extensive future analyses in animal models with endogenous Aβ accumulation.

## Material and methods

### Mice, vertebrate animal procedures

Rodent experimentation was performed with institutional animal care and use committee (IACUC) approval from Duke University School of Medicine. A colony of *hCD155*-tg mice (C57BL/6J background), provided generously by S. Koike [Tokyo Metropolitan Institute of Medical Science, Tokyo, Japan (Koike et al., 1991)], was maintained by the Duke University Division of Laboratory Animal Resources (DLAR) Breeding Core. We used mice ∼6-24 weeks or ∼6 months of age; mice of both sexes were used in equal proportion. The animals were kept under BSL2 conditions in the Duke University School of Medicine Cancer Center Isolation Facility (CCIF) as described previously (Ludwig et al., 2025). All intracerebral inoculations, with stereotactic guidance, were performed to target the central striatum (x=2.0mm, y=3.6mm, z=-0.5mm) with rigorously optimized methods described before (Yang et al., 2023). For tumor cell implantation, 1 x 10^5^ CT2A^hCD155^ cells were inoculated in 5 μl volume at a rate of 2.5 μl/min. All intracerebral virus/mock treatments were performed with a volume of 5 μl at a rate of 2.0 μl/min administering a dose of 5 x 10^7^ pfu [suspended in Dulbecco’s minimal essential medium (DMEM, Invitrogen #11965092)]; mock intracerebral inoculation was done with vehicle (DMEM). Single-dose PVSRIPO was administered on day 7 after tumor implantation; tandem-repeat virus injection occurred on days 7 and 11. Aβ(1-42) HiLyte AF488/647 were administered at a concentration of 2.5 μg in 5 μl [suspended in phosphate-buffered saline (PBS)] using the same stereotactic infusion procedures as described above. For the assay shown in Figure 8A, Aβ HiLyte AF488 was pre-aggregated for 7 days at 37°C prior to implantation. Polio immunization of *hCD155*-tg mice was performed as described previously (Brown et al., 2023). Briefly, mice received two intramuscular injections of PVSRIPO at days-45 and-31 (prior to tumor implantation). The virus inoculum was diluted in PBS to a concentration of 2 x 10^7^ pfu/ml and mixed 1:1 with alhydrogel (Invivogen #vac-alu-50). One 50 μl injection of this mixture was given in each hind quad at each immunization interval.

### Plx5622 and sham diets

CSF1R-inhibitor (Plx5622) treatment and control diets were formulated into AIN-76A rodent chow by Research Diets, Inc. according to a protocol provided by Kim N. Green, Ph.D. (University of California, Irvine). The treatment diet was prepared to contain 1200 ppm Plx5622 (600mg Plx5622 per 500g chow). Plx5622 (MedChemExpress, #HY-114153) was dissolved in 3.6 ml anhydrous DMSO (Sigma-Aldrich, #276855-100ML) by gentle mixing with occasional swirling (no vortexing) until fully solubilized. Poloxamer 407 (125mg; MilliporeSigma, #16758-250G) was subsequently dissolved at room temperature in both Plx5622-containing and vehicle DMSO solutions. The sham diet was identical to the Plx5622 diet except for the absence of active drug. All solutions were protected from light until incorporation into chow. Final diet formulation was performed by Research Diets, Inc. according to their standard protocol.

### Virus, cells, Aβ reagents

PVSRIPO was propagated, purified, and quantified as reported previously (Dobrikov et al., 2022). We used the syngeneic, immunocompetent mouse CT2A glioma cell line, generated by intracerebral implantation of 20-methylcholanthrene containing pellets (Zimmerman and Arnold, 1941), to model malignant glioma in mice. CT2A (RRID:CVCL_ZJ44, a gift from P. Fecci, then Duke University School of Medicine, Durham, NC, USA) was authenticated by whole-exome sequencing. CT2A cells were transduced with hCD155 as described previously to generate the CT2A^hCD155^ model (Ludwig et al., 2025); CT2A^mCherry^ cells were generated in a similar process with a lentiviral vector expressing mCherry (under control of an EF1a promoter) and a puromycin resistance cassette (SignaGen Laboratories #SL100273) according to the manufacturer’s protocol. Briefly, 8 x 10^5^ cells were seeded in 60 mm plates and transduced the following day with the lentiviral construct (MOI 10) with polybrene (5 μg/ml; Millipore Sigma #TR-1003-G) added to culture medium. After transduction (72h), puromycin was added to the culture (2.5 μg/ml) and maintained until cells were of sufficient quantity for sorting. High expressing mCherry^+^ cells were FACS-sorted to establish the CT2A^mCherry^ cell line. CT2A^hCD155^/CT2A^mCherry^ cells were prepared for implantation into *hCD155*-tg mice by propagation in DMEM with 10% fetal bovine serum (FBS; Sigma, #F0926). Cells were harvested when the cultures reached ∼60-70% confluency. Human M059J GBM cells (RRID:CVCL_0400, a gift from M. Bowie, Duke University School of Medicine, Durham, NC, USA) and mouse B16.F10^OVA^ melanoma cells [see information provided in (Brown et al., 2017) and (Ludwig et al., 2025)] were used exclusively as (UV-inactivated) bait for *in vitro* mouse or human microglia phagocytosis assays, respectively (see below). Stock samples of all cell lines were confirmed mycoplasma-free with the MycoStrip Mycoplasma detection kit (Invivogen #rep-mys-10) prior to long-term storage in liquid nitrogen vapor phase. Aliquoted vials of cells were removed from storage and maintained in culture for maximally 4 passages. Aβ (1-42) HiLyte Fluor 488 and 647-labeled Aβ (1-42) (Anaspec #AS-60479-01 and #AS-64161, respectively) were resuspended in 50 μl NH_4_OH and stored in aliquots of 0.2 mg/ml in PBS at-80°C until use.

### Microglia morphology analysis

Tissue sections stained for Tmem119 were imaged at 20x magnification at a resolution of 1024 x 1024, 2048 x 2048, and 4096 x 4096 (see **Microscopy** below). Using ImageJ.tiff images were created of each fluorescent channel and processed for the analysis of morphological features in MicrogliaMorphology/MicrogliaMorphologyR as described previously (Kim et al., 2024). The ImageJ macro MicrogliaMorphology was used as described in Kim *et al*. (Kim et al., 2024) to account for variability in image preparation and acquisition; the code is publicly available on: github.com/ ciernialab/MicrogliaMorphology and /MicrogliaMorphologyR. Optimal local thresholding methods and radius values were selected to accurately segment individual microglial cell fragments or overlapping cells. These parameters were then applied within the macro to guide all downstream morphological analyses. Images were then binarized and converted to greyscale. The resulting images informed individual ROI manager functions to generate depictions of every individual microglial cell, allowing for the assessment of morphology measures. The cells were skeletonized to analyze distinct, specific morphology features. Fractal analysis was performed to measure additional morphology characteristics. MicrogliaMorphologyR wraps multiple packages including tidyverse, Hmisc, pheatmap, factoextra, ImerTest, Ime4, Matrix, SciViews, ggpubr, glmmTMB, DHARMa, rstatix, and gridExtra. Statistical analyses were conducted on GraphPad Prism.

### Microglia isolation, culture, phagocytosis and tumor antigen cross presentation assays

Microglia were isolated from freshly dissected mouse brains or from epilepsy surgery human brain tissues using a CD11b^+^-bead isolation kit (Miltenyi MicroBeads, human & mouse; #130-097-142) as described in detail previously (Bordt et al., 2020). Mouse brain tissue was harvested after transcardiac perfusion with ice-cold sterile PBS. Human brain tissues were de-identified, discarded surplus surgical samples collected under an Institutional Review Board (IRB) declared-exempt protocol. Human/mouse brain tissues of interest (∼80-200 mg fragments) were prepared by mincing with razor blades at 4°C before transfer into 5 ml of Enzyme Digestion Mix [Hanks balanced salt solution (HBSS) without calcium/magnesium, 5% FBS, 10 mM HEPES, 2 mg/ml Collagenase A (Millipore Sigma #11088793001), 28 U/ml DNase I (Millipore Sigma #10104159001)] at 4°C. Samples were transferred for incubation at 37°C (15 min) with agitation (every 5 min), followed by dissociation into single-cell suspensions via sequential trituration through flame-polished Pasteur pipets of decreasing bore diameter (3 steps, 20 passes each), with 15 min interim incubations at 37°C between steps (Bordt et al., 2020). The resulting samples were filtered (70 µm nylon mesh; Sefar #03-70/33) into fresh 15 ml tubes and suspended in HBSS (with calcium/magnesium), then centrifuged at 300 x g (10 min, 4°C). The samples were cleared of Debris/myelin with Debris Removal Solution (Miltenyi Biotec #130-109-398) per manufacturer instructions, and a rinsing spin at 1,000 x g (10 min, 4°C). The pellet was resuspended in MACS Buffer (1 x PBS, 1 mM EDTA, 1% BSA) containing CD11b^+^ MicroBeads (Miltenyi Biotec #130-093-634; 10 µl beads per 90 µl buffer) and incubated at 4°C (15 min). After rinsing cells with centrifugation at 300 x g (10 min, 4°C), CD11b^+^ cells were isolated via LS magnetic columns (Miltenyi #130-042-401) mounted on a MACS Separator (Miltenyi #130-090-976). Flow-through CD11b^−^ cells were collected, and CD11b^+^ cells were eluted by removal of the column from the magnet and plunging twice with 5 ml MACS buffer. CD11b^-/+^ fractions were washed at 300 x g (10 min, 4°C) and resuspended in media selected for the various downstream applications described in this report. For *in vitro* phagocytosis assays, wild-type CT2A, M059J, and B16^OVA^ cells were stained with Cell Trace Violet (Thermo Fisher #C34571) following the manufacturer’s instructions. For microscopic imaging analyses of phagocytosing microglia, Cell Trace Green (Thermo Fisher #C7025) labeling of CT2A cells was used following the manufacturer’s instructions. After the tracer staining procedure, cells (2.5 x 10^6^) were incubated at 37°C (20 min), washed and resuspended with 10ml of DMEM + 10% fetal bovine serum (FBS; 10 min) in a 10 cm culture dish. They were then irradiated by 1hr exposure to ultraviolet light in a biosafety cabinet, confirmed non-viable by trypan blue staining, and stored at 4°C until use.

Microglia were plated at 70% confluency and infected with PVSRIPO (MOI 10) or mock (DMEM), recombinant mouse IFNγ (200 U/ml; R&D Systems, #485-MI), or poly(I:C) (10 μg/ml; Invivogen, high molecular weight polyinosine:polycytidylic acid; LyoVec^TM^ complexed, #tlrl-piclv) and incubated with 200,000 UV-irradiated cells for the duration of the intervals shown in the figures. For cross presentation assays anti-mouse H-2K^b^:SIINFEKL-PE (BioLegend; PE/DazzleTM 594 clone 25-D1.16, #141604; RRID_AB_10895905) was used to assess tumor antigen (SIINFEKL) cross-presentation after B16^OVA^ uptake. At the specified time points, the cultures were rinsed with DMEM to remove non-adherent cells or spun down if microglia were not adherent. For testing microglia phagocytosis of Aβ HiLyte *in vitro*, the material was pre-aggregated as described above, added to microglia at a concentration of 1 μM (96h) and infected with PVSRIPO (MOI 10). For phagocytosis assays with Aβ HiLyte and CT2A^mCherry^ *in vivo*, CD11b^+^-bead isolated microglia were processed into a single-cell suspensions. Microglia were analyzed for phagocytosis tracers (AF488, mCherry), or after staining with antibody probes (see below) by flow cytometry.

### Flow cytometric analyses of mouse and human microglia

We followed flow cytometry procedures outlined in detail previously (Ludwig et al., 2025). Briefly, human/mouse microglia samples were rinsed with, and incubated (20°C, 30 min) in FACS buffer (2% FBS in PBS) with 1% mouse (BioLegend #101319, RRID:AB_1574973) or human (BioLegend #422301, RRID:AB_2818986) TruStain FcX FC receptor blocking solution. 7-AAD viability dye (Thermo Fisher #00-6993-50) was used to stain non-viable cells in samples. Antibodies or isotype controls used for flow cytometry analyses were then added to the suspensions (see Table S2 for a list of all antibodies used) and incubated (20°C, 1h) while shaking (every 15 min) in the dark. After rinsing the cells (1 ml FACS buffer) and centrifugation (1500g, 5 min), the resulting pellet was resuspended in 300 μl FACS buffer in preparation for flow cytometry using the SpectroFlo platform on either Cytek Aurora (Cytek Biosciences) or LSRFortessa (BD Biosciences) instruments.

Microglia were stained for CD45, Cx3cr1, Cd11b, MERTK, and Tmem119 for identification, and a range of additional probes for molecular and functional analyses (see Table S2). For intracellular vRNA staining, HCR-FISH was combined with flow cytometry in procedures and with hybridization probes that have been described in detail previously (Ludwig et al., 2025). Positive vs. negative staining, appropriate compensation/spectral unmixing, and specificity of antibody staining was defined with the use of confirmed negative cell populations, single stain controls and isotype controls using beads and unstained microglia as reported before (Ludwig et al., 2025). VersaComp beads (Beckmann-Coulter) and staining of fresh cells were used to standardize compensation; flow cytometry data were analyzed with FlowJo v10.10 (BD Biosciences).

### RT-qPCR, HCR-FISH

RT-qPCR analyses were conducted on a QuantStudio3 (Thermo Fisher) platform; template abundance was determined using the 2^_ΔΔCT method, as described in detail previously (Ludwig et al., 2025). For RT-qPCR analyses from microglia harvested from PVSRIPO-treated brains, mice were perfused with ice-cold PBS, their brains were harvested, split at the midline using a razor blade for experiments comparing ipsilateral vs. contralateral vRNA burden, and processed for CD11b^+^-bead microglia isolation (see above). Total RNA was isolated from microglia harvested from PVSRIPO-treated brains, or from microglia infected *in vitro*, by Trizol (Invitrogen) extraction. RNA isolation by Trizol-chloroform extraction, column purification (GeneJet RNA purification; Thermo Fisher #K0731) and determination of RNA concentration with a nanodrop were carried out as described previously (Ludwig et al., 2025). For RT-qPCR analysis, RNA samples were subjected to DNase removal with PrimeScript™ FAST RT Reagent Kit (Takara Bio) and reverse transcribed into cDNA. Viral (+)strand RNA was analyzed with custom primers for PVSRIPO (Dobrikova et al., 2012), as described before (Ludwig et al., 2025). One-step qPCR was performed with standard primer pairs for MDA5 (#Hs00223420_m1 FAM), IRF7 (#Hs01014809_g1 FAM), and TREM2 (#Hs00219132_m1 FAM) (all Applied Biosystems) using the TaqMan Fast Advanced Master Mix (Applied Biosystems).

### Microscopy

For immunohistochemistry/immunofluorescence (IF) assays mice were perfused with ice-cold PBS. Brain and deep cLN tissues were dissected, embedded in OCT medium (TissueTek, Sakura #4583), flash-frozen over liquid nitrogen, and stored at-80°C until cryosectioned and fixed in 4% paraformaldehyde (15 min). The sections were permeabilized (0.05% Triton X-100 in PBS-T), incubated in blocking buffer (Donkey or Goat Serum, 10%; Sigma-Aldrich #9663 and Thermo Fisher #31872, respectively) before incubation with primary antibody (1:200 dilution; 24h, 4°C) (antibody probes used are listed in Table S2). After staining with primary antibody, the sections were rinsed (3x, TBS; Invitrogen #J77500.K2) and incubated with correlating secondary anti-species antibodies: i) goat anti-rabbit IgG (H+L) (Invitrogen #A32733TR, RRID:AB_2866492); ii) donkey anti-mouse IgG (H+L) AF594 (Invitrogen #A-21203, RRID:AB_2535789); or iii) goat anti-rabbit IgG (H+L) AF488 (Invitrogen #A32731, RRID:AB_263280) at 20°C (2h). Sections were counterstained with Hoechst 33342 (Thermo Fisher #62249) and mounted using VectaShield anti-fading medium (Vector Labs #H-1000-10). HCR-FISH for imaging the spatial distribution of PVSRIPO (-)strand vRNA was carried out with methods and molecular probes described previously (Ludwig et al., 2025). For imaging of cultured cells, microglia were cultured on poly-L-lysine 12 mm coated coverslips (Corning #354085). The cell preparations were processed for IF as described for cryosections above; methods and molecular probes used in HCR-FISH of (-)strand vRNA were described in detail previously (Ludwig et al., 2025). Microscopic images were acquired on a Leica Stellaris 8 scanning confocal microscope at the Light Microscopy Core Facility (Duke University) and processed using Fiji(ImageJ) as reported before (Ludwig et al., 2025).

### Single cell RNAseq

GBM *ex vivo* sclice culture assays were performed as reported in detail previously (Brown et al., 2021). Samples were snap-frozen and sent to Azenta Life Sciences (Genewiz) for nuclei isolation, library preparation, sequencing, and alignment. Libraries were made using the Chromium Single Cell 3’ Gene Expression v4 platform aiming for 10,000 nuclei per sample and 50,000 reads per nucleus. Sequencing was performed on an Illumina NovaSeq X Plus (2 x 150 bp). Sequencing reads were processed using the Cell Ranger pipeline (v8.0.0; 10x Genomics). Gene expression feature matrices were generated using the cellranger count function with default parameters unless otherwise specified. Reads were aligned to the human reference transcriptome (GRCh38, refdata-gex-GRCh38-2020-A) with read lengths of 28 bp for read 1 and 98 bp for read 2, chemistry automatically detected, and intronic reads included during quantification. Further analyses were performed using the filtered feature matrices produced by Cell Ranger in R (4.5.2). Analyses were conducted primarily using the Seurat package (v5.3.1), and functions from this package were used unless otherwise stated. Other packages used were scuttle (v1.18.0), ggplot2 (v4.0.1), msigdbr (v25.1.1), fgsea (v1.34.2). Data were imported using ‘Read10x_h5()’ and ‘CreateSeuratObject()’. Cells were filtered for quality based on RNA counts (nCount_RNA), number of detected genes (nFeature_RNA), and percent of mitochondrial RNA (defined as genes beginning with “MT-“).

Outliers were identified using scuttle’s ‘isOutlier()’ (min.diff = 0.5). Outliers were detected in both directions for features and counts and only in the upper direction for mitochondrial RNA. Highly variable features were selected using ‘FindVariableFeatures()’ (selection.method = “vst”, nfeatures = 2000). Data were then normalized using ‘NormalizeData()’, and samples were integrated to correct for treatment effects for cell-type annotation using the ‘SelectIntegrationFeatures()’, ‘FindIntegrationAnchors()’, ‘IntegrateData()’, and ‘JoinLayers()’. Dimension reduction and clustering were performed using ‘RunPCA()’, ‘FindNeighbors()’ (dims = 1:15), ‘FindClusters()’ (resolution = 0.75), ‘RunUMAP()’, and ‘RunTSNE()’. Cluster markers were identified using ‘FindAllMarkers()’. Clusters displaying non-specific markers indicative of low quality were removed and clustering repeated. Microglia and tumor-associated myeloid cells were identified at this stage, while non-immune cells were grouped together based on their UMAP clustering patterns. The resulting labels were then applied to the non-integrated data, and clustering and visualization were performed using the same workflow. UMAP visualization was generated using ‘DimPlot()’. Scaled counts were generated using the ‘ScaleData()’. Microglia were subsequently subsetted using ‘subset()’, and genes of interest in microglia were displayed using ggplot2’s ‘ggplot()’, ‘geom_violin()’, ‘geom_jitter()’, and ‘facet_wrap()’. Differential expression was performed using ‘FindMarkers()’ which implements a Wilcoxon rank-sum test with Bonferroni correction for multiple comparisons for adjusted p-values. Hallmark pathways were accessed from the Molecular Signatures Database (MSigDB) using msigdbr’s ‘msigdbr()’ (species = “Homo sapiens”, collection = “H”). The set of ISGs was downloaded from (Yu et al., 2025). GSEA was performed on ordered differentially expressed genes from ‘FindMarkers()’ using fgsea’s ‘fgsea()’ with 10,000 permutations (eps = 0.0, minsize = 15, and maxSize = 500). For hallmark pathways, were collapsed using fgsea’s ‘collapsePathways()’ (pval.threshold = 0.05, gseaParam = 1) to remove highly overlapping sets. Enrichment statistics were calculated using fgsea’s implementation of a Kolmogorov-Smirnov-like statistic with p-values derived from permutation testing and adjusted using Benjamini-Hochberg false discovery rate (FDR) procedure. Results are displayed using fgsea’s ‘plotGseaTable()’ with gseaParam = 1. P2RY12 and GPR34 were selected (as microglia distinguishing markers and displayed using ‘FeaturePlot()’. Genes were subsetted on type-I IFNs with any detected transcripts across both samples where percent-positive cells were defined as the proportion of cells with at least one transcript of the given gene to total population.

## Statistics

Statistical analyses of the reported data were conducted with R and GraphPad. Specific methods employed, as indicated, are described in the corresponding figure legends. Error bars = SEM.

## Acknowledgments

We thank Arilyn Lynch, Merrie Burnett, Elizabeth Thomas, Daryl Walker and Dianne Satterfield of the Preston Robert Tisch Brain Tumor Center Tissue BioRepository at Duke University School of Medicine for their help with human tissue collection and logistics, and mouse tissue histological processing and cryosectioning.

This work was supported by PHS grants R01 NS108773 (M.G.), R01 CA281320 (M.G.), and R00 CA263021 (M.B.). We acknowledge an NIH instrumentation grant for the Leica Stellaris confocal microscope used in our studies at the Duke University Light Microscopy Core Facility (1S10OD034340-01A1).

## Author contributions

mouse glioma model studies, G.C. Carter, and M. Gromeier; Plx5622 depletion studies, G.C. Carter, Z.P. McKay, and M. Gromeier; morphology analyses, G.P. Carter; microglia isolation, G.P. Carter; HCR-FISH, G.P. Carter; flow cytometry, G.P. Carter and M.A. Katz; IF, G.C. Carter, L. Disla and M.A. Katz; imaging and microscopy, G.P. Carter, L. Disla, and M.A. Katz; scRNAseq, G.P. Carter, Z.P. McKay and M.C. Brown; phagocytosis assays, G.P. Carter, D.N. White, and M.A. Katz; RT-qPCR, G.P. Carter; injectable Aβ HiLyte mouse model, G.P. Carter and M. Gromeier; human tissue procurement, G.P. Carter and D. Southwell; manuscript writing and editing, G.P. Carter, M.C. Brown, and M. Gromeier. All authors reviewed and edited the manuscript.

