## Supplemental Figure 1, Table 2 for "Viral Microglia Reprogramming Clears Oligomeric Neurotoxic Debris"

**SUPPLEMENTAL MATERIAL**

Carter *et al*.
MDA5-mediated microglia reprogramming

### Viral Microglia Reprogramming Clears Oligomeric Neurotoxic Debris

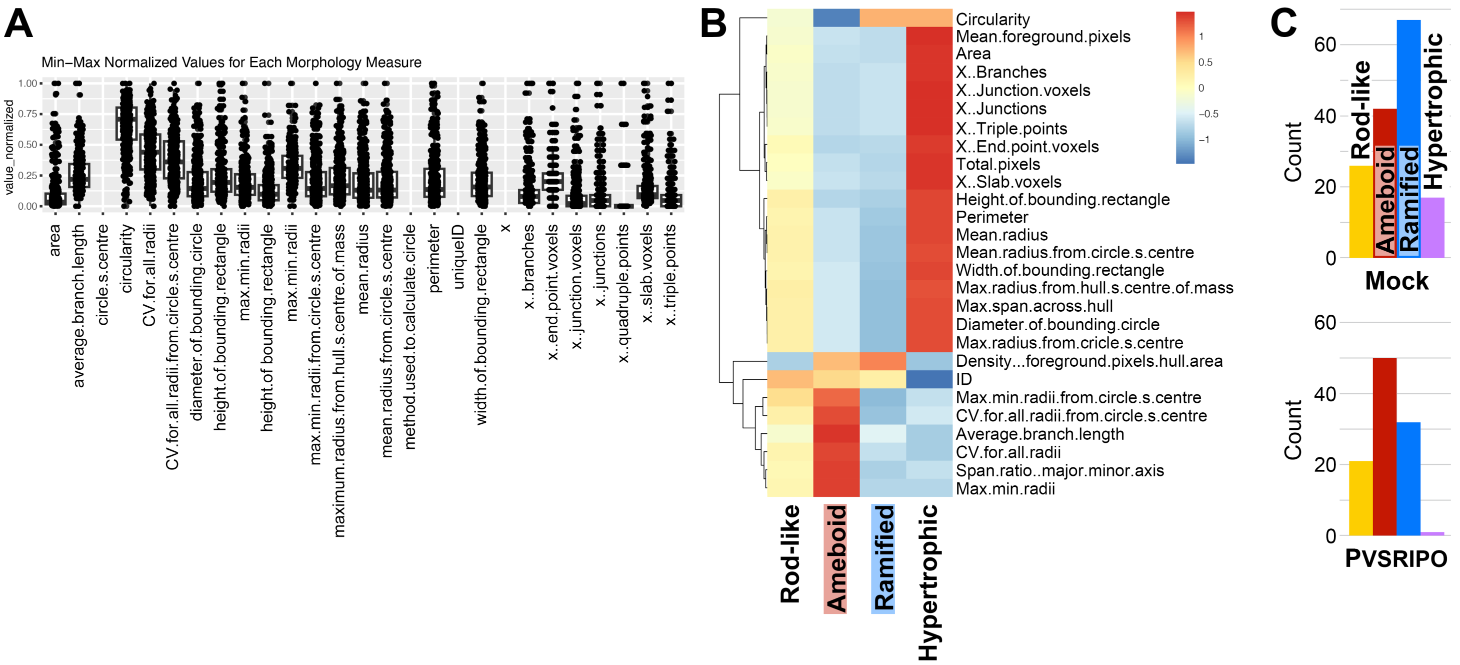


**Figure S1 (Related to Figure 1C).** Validating the microglia morphology analyses tool reported by Kim *et al*. (2024) for our approach. Extended data from analyses of a set of “training” images stained for Iba1 using MicrogliaMorphology (ImageJ) and MicrogliaMorphologyR (R package) toolsets. **(A)** Distribution of 27 distinct morphological features found in cell shapes from Images shown in Figure 1C [ (Mock-treated) glioma-bearing brain, (PVSRIPO-treated) glioma-bearing brain were combined]. **(B)** MicrogliaMorphology-assigned clusters based on the defined 27 features of Iba1-stained phagocytes in sections shown in Figure 1C. **(C)** Preliminary analyses of changes in cluster frequency in Mock- vs. PVSRIPO-treated glioma-bearing brain, based on validation experiments with Iba1-stained samples.

**Table S1 (Related to Figures 3 and 4; attached separately).** Conserved markers across treatments for astrocyte (Astrocyte _markers), neoplastic (Cancer_markers), bone marrow-derived myeloid cells (Macrophage_ markers), microglia (Microglia_markers), neurons (Neuron_markers), and pericyte & fibroblast (Pericyte_Fibroblast_markers) clusters; differentially expressed genes conserved across treatment comparing microglia to bone marrow-derived myeloid cells (Microglia_v_ Macrophages _markers); treatment markers within microglia comparing PVSRIPO to mock (RIPO_v_Mock_microglia).

| **Antigen** | **Fluorophore** | **Vendor** | **Cat #** | **Dilution** | **RRID** |
|---|---|---|---|---|---|
| Mouse: | | | | | |
| CD45 | BUV395 | BD | 564279 | 100x | AB_2651134 |
| CD3 | PE | Biolegend | 100206 | 100x | AB_312663 |
| IA/IE | BV785 | Biolegend | 107645 | 100x | AB_2565977 |
| CD11c | APC | Biolegend | 117310 | 100x | AB_313779 |
| CD11c | APC/Cy7 | Biolegend | 117324 | 100x | AB_830649 |
| CD11c | BV421 | Biolegend | 117330 | 100x | AB_11219593 |
| F4/80 | BUV805 | BD | 749282 | 100x | AB_2873657 |
| IA/IE | BV711 | Biolegend | 107643 | 100x | AB_2565976 |
| F4/80 | BV605 | Biolegend | 123133 | 100x | AB_2562305 |
| CD11b | BV711 | Biolegend | 101236 | 100x | AB_11203704 |
| Ly6C | PerCP-Cy5.5 | BD | 560525 | 100x | AB_1727558 |
| F4/80 | PE-Cy5 | Biolegend | 123112 | 100x | AB_893482 |
| Ly6G | BV605 | Biolegend | 127639 | 100x | AB_2565880 |
| Siglec-F | PE-Cy7 | Biolegend | 155528 | 100x | AB_2890715 |
| CD3 | BV605 | Biolegend | 317322 | 100x | AB_2561911 |
| CD4 | FITC | Biolegend | 100406 | 100x | AB_312691 |
| H-2KB/D | PE | Biolegend | 114608 | 100x | AB_313599 |
| CD45 | BUV737 | Biolegend | 612778 | 100x | AB_2870107 |
| CD86 | FITC | Biolegend | 105006 | 100x | AB_313149 |
| CD68 | APC/Fire-750 | Biolegend | 137042 | 100x | AB_2910295 |
| H2KB | PE-Cy7 | Invitrogen | 25-5958-82 | 100x | AB_2573505 |
| Cx3cr1 | PerCP-Cy5.5 | Biolegend | 149010 | 100x | AB_2564494 |
| CD80 | PE-Cy7 | Biolegend | 104734 | 100x | AB_2563113 |
| Ki-67 | BV421 | Biolegend | 652411 | 100x | AB_2562663 |
| Tmem119 | PE-Cy7 | Invitrogen | 25-6119-82 | 100x | AB_2848312 |
| CD68 | FITC | Biolegend | 137006 | 100x | AB_10578412 |
| TREM2 | FITC | Invitrogen | MA5-28223 | 100x | AB_2745193 |
| CD40 | PE-594 | Biolegend | 124630 | 100x | AB_2572185 |
| H2KB: SIINFEKL | PE | Biolegend | 141604 | 100x | AB_10895905 |

| **Antigen** | **Fluorophore** | **Vendor** | **Cat #** | **Dilution** | **RRID** |
|---|---|---|---|---|---|
| Human: | | | | | |
| CD11b | APC | Biolegend | 982604 | 200x | AB_2632619 |
| CD11c | BUV395 | BD | 563787 | 200x | AB_2744274 |
| CD155 | PE | Biolegend | 337610 | 200x | AB_2174019 |
| HLADR | BV786 | Biolegend | 307642 | 200x | AB_2563461 |
| CD68 | APC/Fire-750 | Biolegend | 333824 | 200x | AB_2800878 |
| Tmem119 | Alexa 647 | Biolegend | 853312 | 200x | AB_3097501 |
| Cx3cr1 | BV785 | Biolegend | 341628 | 200x | AB_2810535 |
| MERTK | PE-Cy7 | Biolegend | 367610 | 200x | AB_2687287 |
| IF (mouse): | | | | | |
| Iba1 | | CST | 17198 | | AB_2820254 |
| Tmem119 | | CST (E3E10) | 90840 | | AB_2928137 |
| ISG15 | | Invitrogen | 703132 | | AB_2784563 |
| A11 | | Thermo F. | AHB0052 | | AB_10376183 |

**Table S2.** Antibodies used in this study.
